# Programmable and Switchable RNA Scaffolds for Synthetic Condensate Engineering in Mammalian Cells

**DOI:** 10.64898/2026.02.14.705909

**Authors:** Zhaolin Xue, Oyeshik Mukherjee, Lan Mi, Yizhan Guo, Ru Zheng, Chang Yuan, Mingxu You

## Abstract

The ability to engineer synthetic biomolecular condensates in living cells offers new opportunities to control intracellular organization, yet robust and programmable RNA-based systems have remained limited. Here, we introduce genetically encoded, modular platforms that generate RNA-driven condensates using nanostar-derived scaffolds. Systematic comparison of repeat-based and *de novo* designs identified nanostar variants that reliably assemble nuclear condensates in mammalian cells. Unexpectedly, condensate formation in cells is governed primarily by double-stranded RNA stems that recruit endogenous RNA-binding proteins, rather than by the kissing-loop interactions that drive assembly *in vitro*. This mechanistic shift highlights the divergence between cellular and *in vitro* environments and accounts for the limited orthogonality among scaffolds. Sequence refinement to reduce nonspecific pairing improved homotypic assembly and enhanced orthogonality. We further demonstrated functional compartmentalization by recruiting protein and RNA clients to modulate their stability and activity, and we incorporated an acyclovir-responsive allosteric switch to achieve reversible, small-molecule control of condensation. Together, this work establishes a versatile RNA-based toolkit for constructing programmable cellular compartments, advancing strategies for controlling RNA–protein organization and enabling new biosensing and therapeutic applications.

## INTRODUCTION

Cellular function relies on highly organized spatiotemporal regulation, exemplified by biomolecular condensates. These membraneless organelles, formed through liquid–liquid phase separation and related percolation process, compartmentalize biochemical activities to regulate gene expression, signaling, and stress responses^1^. Their dynamic assembly and disassembly are essential for cellular organization and adaptability, and disruption in these processes are linked to diseases such as neurodegeneration and cancer^2,3^.

RNA plays a central role in many natural condensates. Beyond its genetic function, RNA acts as both a structural scaffold and a regulatory client, promoting phase separation through programmable, multivalent interactions including base pairing, G-quadruplex formation, and loop–loop contacts^4,5^. This inherent programmability makes RNA a powerful platform for engineering synthetic biological systems. However, most synthetic condensate designs continue to rely on protein-based scaffolds built from disordered polypeptides or modular protein domains^6,7^. In contrast, RNA-driven condensates offer unique advantages^2,8^, yet current approaches do not fully leverage RNA’s structural diversity, tunability, and natural integration into cellular RNA-processing pathways that enable precise control over RNA-centric biology.

A major gap in the field is the absence of robust, generalizable platforms for constructing RNA-driven synthetic condensates in mammalian cells, particularly those support orthogonal, dynamic, and functional control. Existing systems predominantly target proteins and are difficult to adapt for RNA-specific functions. Developing tools to synthetically control RNA condensation would unlock new avenues to manipulate RNA localization, stability, translation, and interaction networks, directly engaging a fundamental layer of cellular regulation.

Here, we address this gap by introducing a modular, genetically encodable platform for engineering programmable RNA-driven condensates in mammalian cells. We designed a series of circularized RNA scaffolds, derived from *de novo* nanostar motifs^9,10^, that robustly form condensates in cells. Notably, their assembly is governed by an unexpected mechanism involving double-stranded RNA stems. This platform supports selective recruitment and functional modulation of protein and RNA clients, improves orthogonality through sequence design, and incorporates a small-molecule-responsive allosteric switch for reversible control. By harnessing RNA’s dual role as an informational polymer and a structural scaffold, this system provides a versatile and programmable framework for spatial and functional engineering in mammalian cells, with board applications in synthetic biology and therapeutic design.

## RESULTS

### Evaluating repeat-based and *de novo* RNA motifs for condensate assembly

To engineer RNA condensate scaffolds, we first compared a set of RNA motifs for their ability to drive condensation *in vitro*. Although several RNA motifs can form condensates, their relative condensation strength, particularly in cellular environments, remain unclear. For systematic comparison, we synthesized fluorogenic aptamer-tagged RNA strands containing 47×CAG^11–13^, 47×CUG^14^ or 29×G_4_C_2_^11,15^ repeats. We also prepared *de novo-*designed RNA motifs, including 5×Corn, 5×Beetroot^16^, and the “design B” (Bro-NS) and “design A” (Pep-NS) RNA nanostars^10^ (Supplementary Fig. 1, Table 1).

When annealed in 4 μM RNA, 20 mM MgCl_2_, and added the cognate fluorophores, all motifs formed condensates (Fig. 1a). The repeat-based motifs generated spherical condensates with diameters of ∼1.1–1.8 μm, whereas 5×Corn and 5×Beetroot produced droplet networks of ∼1.4 μm (Fig. 1b). Among them, 5×Corn formed the most extended network, exhibiting the highest aspect ratio (Fig. 1c). Partition-ratio measurement showed that the repeat motifs displayed ∼4.5–8.5 enrichment, while the *de novo* motifs showed ∼2–4 (Fig. 1d). Notably, the 2×dimerized Broccoli tag (2dB)^17^ increased the partition ratio of 47×CAG condensates without altering their size, likely due to G-quadruplex-mediated RNA packing and enhanced multievent interactions.

**Fig. 1.**
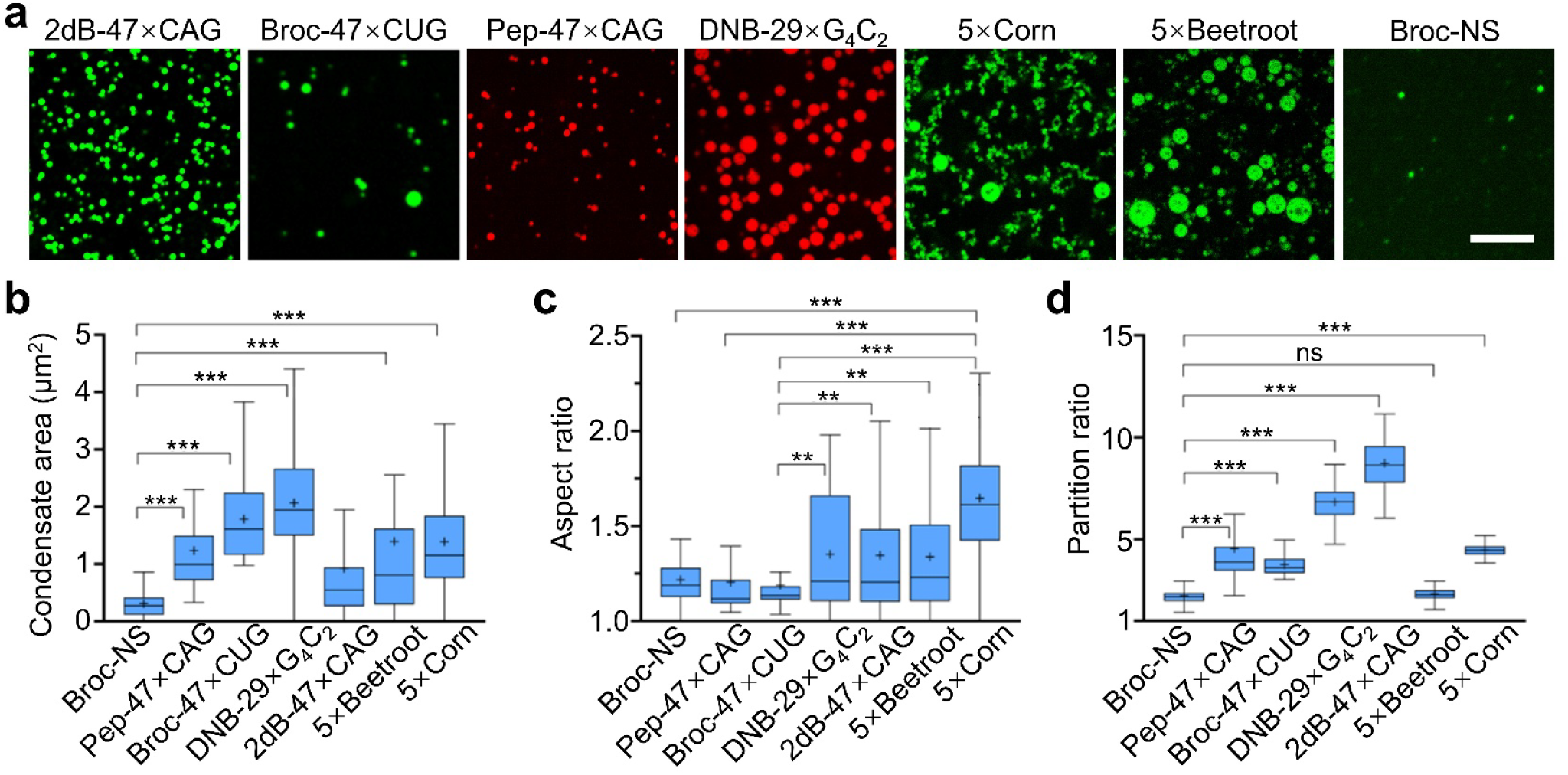
RNA scaffolds induce condensation *in vitro*. **a**, Confocal fluorescence images of RNA condensates formed by repeat-based scaffolds (47×CAG, 47×CUG, 29×G_4_C_2_) and *de novo-*designed scaffolds (5×Corn, 5×Beetroot, nanostar [NS]). Fluorogenic RNA tags—2d×Broccoli (2dB), Broccoli (Broc), Pepper (Pep), dinitroaniline-binding aptamer (DNB), or intrinsic Corn and Beetroot fluorescence—were used for visualization. Samples contained 4 μM RNA and 20 mM MgCl_2_, with dyes added as appropriate (80 μM DFHBI-1T, 2 μM HBC620, 0.5 μM TMR-DN, or 2 μM DFHO). RNAs were annealed at 95 °C for 2 min and cooled to 37 °C at –0.5 °C/min. The dyes were added for a further 30 min incubation at 37 °C before imaging at room temperature. Scale bar: 10 μm. **b**–**d**, Quantification of condensate area, aspect ratio, and partition ratio from **a**, displayed as Tukey box-and-whisker plots. Boxes represent the 25th and 75th percentiles; central lines indicate the medians; whiskers extend to 1.5× the interquartile range; means are indicated by “+”. For each scaffold, 75 condensates from ≥3 independent replicates were analyzed. Scaffolds are ordered by length: Broc-NS (257 nt), Pep-47×CAG (299 nt), Broc-47×CUG (304 nt), DNB-29×G_4_C_2_ (414 nt), 2dB-47×CAG (463 nt), 5×Beetroot (1156 nt), 5×Corn (1181 nt). Statistical significance was assessed by one-way ANOVA with Tukey’s post-hoc test (\*\*\**p* ≤ 0.001; \*\**p* ≤ 0.01; \**p* ≤ 0.05; ns, *p* > 0.05).

The Broc-NS motif produced relatively small condensates (∼0.6 μm), differing from previously reported co-transcriptional NS condensates. This reduced size likely reflects annealing-induced stem– stem interactions between NS arms, which disrupt multivalent kissing-loop interactions and limit condensation^10,18^. To restore NS condensation, we applied a melt-and-hold protocol^10^ in which RNA strands were heated at 70 °C, rapidly cooled to 37 °C, and incubated at 37 °C for 6 or 24 hours. This treatment markedly increased condensate size from ∼3.6 to ∼6.4 μm and raised the partition ratio to ∼10 (Supplementary Fig. 1). Overall, RNA multivalency, stacking, and crosslinking strength collectively influence condensate formation, and isothermal incubation generally enhances NS condensate growth.

### Genetically encoded RNA scaffolds for condensate formation in mammalian cells

We next asked whether these RNA motifs could serve as genetically encoded scaffolds to drive RNA condensation in mammalian cells. To achieve high RNA expression, we used the pAVU6+27-Tornado vector^19^ to encode each circularized RNA motif fused to its fluorogenic RNA aptamer. The F30 three-way junction^20^ was included to ensure proper aptamer folding and to stabilize the stem required for Tornado circularization. The repeat-based motifs—47×CAG, 47×CUG, and 17×G_4_C_2_—were fused to the F30 3’-arm, with the corresponding fluorogenic aptamer placed on the 5’-arm, generating c2dB-47×CAG, cBroc-47×CUG, and cDNB-17×G_4_C_2_, where “c” denotes circularized RNA (Fig. 2a). The 5×Corn and 5×Beetroot constructs were used directly, with Pepper fused to 5×Beetroot. The Broc-NS motif was truncated at the 3’-end and fused to the F30 5’-stem for Tornado circularization while preserving its three kissing loops, yielding cNS3 (Fig. 2a).

**Fig. 2.**
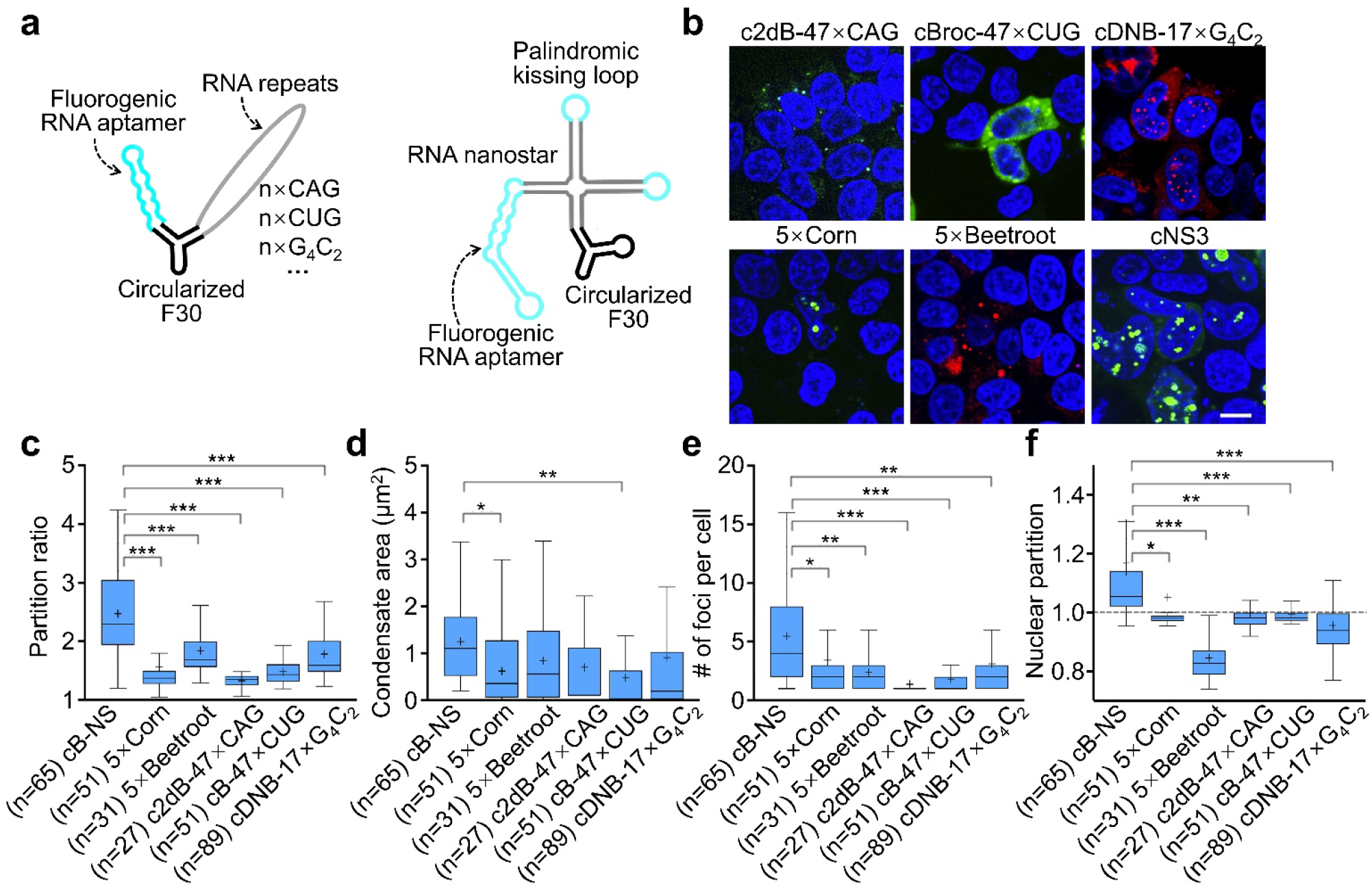
Engineering RNA scaffolds for cellular condensate formation. **a**, Schematic of circularized RNA scaffold fused to an F30 three-way junction and expressed from pAVU6+27-Tornado vector for robust production in mammalian cells. The circular RNA nanostar (cNS3) contains three kissing loops and an embedded Broccoli aptamer. Additional fluorogenic RNAs— 2d×Broccoli (2dB), Broccoli (Broc) and DNB—were fused to 47×CAG, 47×CUG, or 17×G_4_C_2_ repeats using a circularized F30 junction. **b**, Confocal fluorescence images of HEK293T cells expressing circularized RNA scaffolds. Previously reported 5×Corn and 5×Beetroot constructs were used directly, with fluorogenic Pepper inserted into 5×Beetroot for visualization. Cells were incubated with the appropriate dyes (40 μM DFHBI-1T, 1 μM TMR-DN, 40 μM DFHO, or 5 μM HBC620) and 5 mM Mg^2+^ for 30 min. Hoechst 33342 was used for nuclear staining. Scale bar: 10 μm. **c**–**d**, Quantification of partition ratio, condensate area, condensate number per cell, and nuclear partition from **b**, displayed as Tukey box-and-whisker plots. Boxes represent the 25th, 50th, and 75th percentiles; whiskers extend to 1.5× the interquartile range; means are shown as “+”. Data represents n cells from ≥3 replicates for each scaffold. Statistical significance was assessed using one-way ANOVA with Tukey’s post-hoc test (\*\*\**p* ≤ 0.001; \*\**p* ≤ 0.01; \**p* ≤ 0.05; ns, *p* > 0.05). Nonsignificant comparisons are omitted from the plots.

All constructs formed condensates to varying degrees following transient transfection in HEK293T cells (Fig. 2b). Among them, cNS3 exhibited the most robust condensation, with a partition ratio of ∼2.4, approximately five condensates per cell, and a slightly larger diameter of ∼1.2 μm (Fig. 2c–e). cNS3 condensates localized predominantly to the nucleus, as indicated by the nuclear-to-cytoplasmic fluorescence ratio (Fig. 2f). In contrast, c2dB-47×CAG produced minimal condensates despite its enhanced condensation *in vitro* (Fig. 2b–f), consistent with reports that CAG repeats undergo lysosomal degradation^21^. Supplementing the imaging medium with 5 mM MgCl_2_ produced no major changes in cNS3 condensates, aside from a modest increase in condensate size (∼1.5 μm) and a slight reduction in partition ratio (Supplementary Fig. 2a–e). Further cell viability test also indicated no cytotoxicity of NS condensates in HEK293T cells (Supplementary Fig. 2f). Together, these results identify cNS3 as a strong scaffold for driving RNA condensation in cells.

We then examined whether cNS3 condensates form in other mammalian cell lines. Transient transfection of cNS3 into HeLa and MCF-7 cells produced similar nuclear condensates as observed in HEK293T cells (Supplementary Fig. 3). Compared with HEK293T cells, HeLa and MCF-7 cells exhibited larger condensates (∼2.1 μm) and a reduced partition ratio (∼1.8), with approximately nine condensates per cell, likely reflecting their larger cell size (Supplementary Fig. 3). Thus, cNS3 consistently forms nuclear condensates across diverse cell types, demonstrating its utility as a generalizable scaffold for RNA condensation in mammalian cells.

### Structural determinants of NS condensate formation in cells

Next, we asked whether the mechanism that governs NS condensation *in vitro* persists in cells. *In vitro*, NS condensate formation is primarily driven by kissing-loop interactions^9,10^. To test whether cNS3 condensation depends on these interactions, we generated cNS2 and cNS0, in which the kissing loops were replaced with scrambled sequences, leaving two or zero active loops, respectively. When characterized *in vitro*, linearized RNA scaffold subjected to the melt-and-hold protocol showed that NS3 formed condensates comparable to the original “design B” NS, whereas NS2 and NS0 failed to form condensates and instead displayed diffuse fluorescence even under 20 mM MgCl_2_ (Supplementary Fig. 4). This agrees with previous observations that at least three kissing loops are required for NS condensation in vitro^10,22,23^.

However, both cNS2 and cNS0 still formed nuclear condensates after transient transfection in mammalian cells (Fig. 3a, Supplementary Fig. 3), indicating that NS condensation in cells is not primarily driven by kissing-loop interactions. We then examined whether specific structural features of the NS scaffold dictate condensate formation in cells. To assess this, we designed three structural variants— ctNS2, csNS3, and cuNS3. ctNS2 contains a truncated (“t”) set of stems with two remaining NS arms, whereas csNS3 and cuNS3 contain shortened stems or an unfolded four-way junction (“s” and “u”), respectively. The final digit in each name (“2” or “3”) denotes the number of active kissing loops.

All these constructs formed nuclear condensates in HEK293T cells (Fig. 3a). ctNS2 exhibited partition ratio, condensate size, number, and nuclear enrichment comparable to cNS3 (Fig. 3b–e). In contrast, csNS3 and cuNS3 showed significantly reduced partition ratios and high cytoplasmic fluorescence, with nuclear partitioning below one (Fig. 3b, e), indicating lower segregation, higher saturation concentration, and reduced condensation driving force. csNS3 produced smaller condensates but in greater number than cNS3 (Fig. 3c, d), likely due to shortened NS arms limiting interaction range^10^, whereas cuNS3 displayed condensate size and number similar to cNS3 (Fig. 3c, d).

**Fig. 3.**
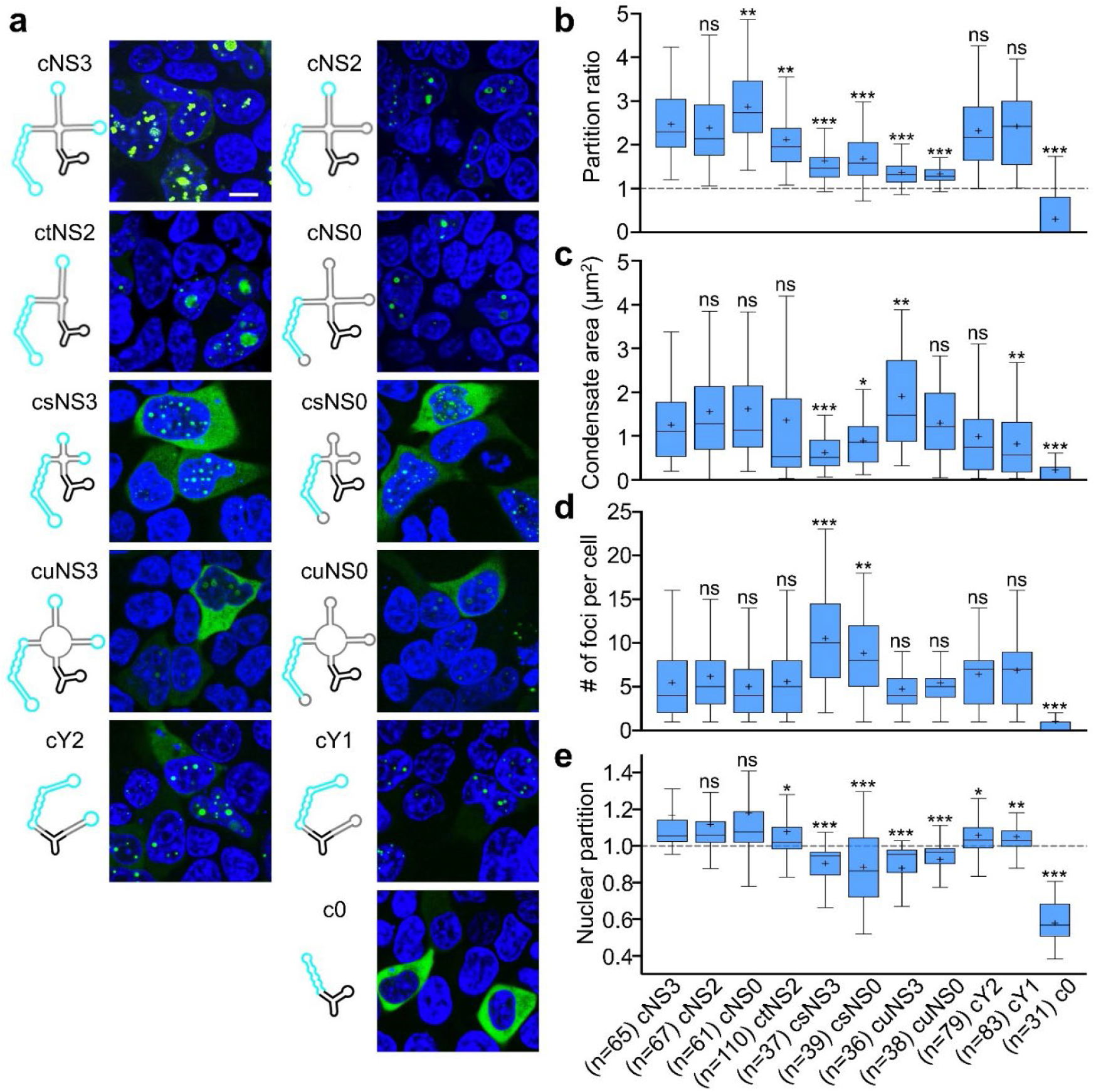
RNA nanostar (NS) variants induce cellular condensate formation via double-stranded RNA stems. **a**, Schematics of circular nanostar (cNS) variants and corresponding confocal fluorescence images in HEK293T cells. Variants with truncated stems, shortened stems, or an unfolded four-way junction are designated ctNS, csNS and cuNS respectively. A Y-shaped variant is denoted cY. The circularized F30 junction is labeled “c”, and the final digit (3, 2, 1, 0) indicates the number of active kissing loops. Live-cell imaging was performed after a 30-min incubation with 40 μM DFHBI-1T and 5 mM Mg^2+^. Hoechst 33342 was used for nuclear staining. Scale bar: 10 μm. **b**–**e**, Quantification of partition ratio, condensate area, condensate number per cell, and nuclear partition from **a**, displayed as Tukey box-and-whisker plots. Boxes represent the 25th, 50th, and 75th percentiles; whiskers extend to 1.5× the interquartile range; means are shown as “+”. Data represents n cells from ≥3 independent replicates. Statistical comparisons were performed using two-tailed Student’s *t*-tests relative to cNS3 (\*\*\**p* ≤ 0.001; \*\**p* ≤ 0.01; \**p* ≤ 0.05; ns, *p* > 0.05).

Scrambling all kissing loops in csNS3 and cuNS3 to generate csNS0 and cuNS0 did not eliminate condensate formation (Fig. 3a), further confirming the minimal role of kissing-loop interactions in cellular NS condensation. Together, these results show that the NS four-way junction is not required for condensation in cells, whereas the integrity of the double-stranded RNA stems is critical, as their shortening or unfolding disrupts condensate formation.

We next examined whether NS RNA duplexes drive condensate formation in cells. To simplify the scaffold, we designed cY2, cY1 and c0. The c0 construct contains only the circularized F30 with Broccoli fused to the F30 5’-arm. For cY2 and cY1, a 17-bp duplex was inserted into the F30 3’-arm and kissing loops were added to both arms, generating a “Y-shaped” scaffold with two or one active kissing loops, respectively. When expressed in HEK293T cells, both cY2 and cY1 formed prominent nuclear condensates with partition ratios, sizes, and condensate numbers comparable to cNS3 (Fig. 3a–e). In contrast, c0, which lacks the inserted duplex, showed only cytoplasmic fluorescence and did not form condensates (Fig. 3a–e). These results indicate that the dsRNA stem is the primary determinant of NS-mediated condensate formation in cells, likely through interaction with nuclear components.

### Identifying nuclear compartments associated with NS condensates

Next, we sought to identify nuclear components that interact with the NS. Because many subnuclear compartments contain RNA-binding proteins (RBPs) that recognize highly structured RNA^24– 28^, and NS condensation in the nucleus depends on RNA duplexes, we examined whether NS structures colocalize with specific nuclear compartments, beginning with the nucleolus^25,29^. To assess potential nucleolar association, we expressed Pepper-tagged NS (P-cNS3) and stained cells with nucleolar-ID. P-cNS3 condensates did not colocalize with nucleolar signals (Fig. 4a, Supplementary Fig. 5a, b). Likewise, cNS3 condensates showed no colocalization with BFP-tagged nucleolin, a nucleolar marker^30^. These results indicate that NS condensates are unlikely to interact with the nucleolus (Fig. 4b, Supplementary Fig. 5c).

**Fig. 4.**
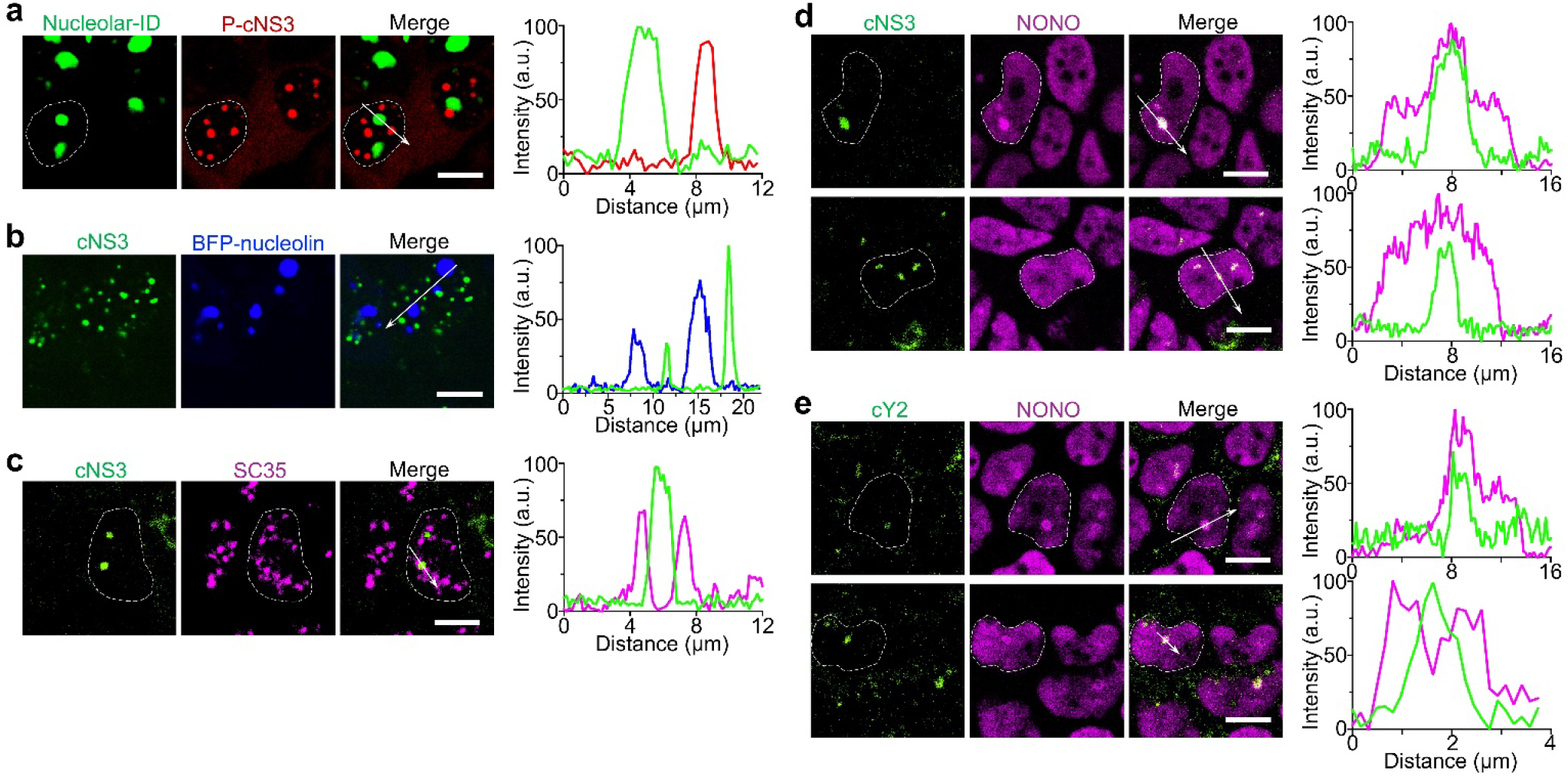
RNA nanostar (NS) condensates colocalize with paraspeckles. **a**, NS condensates are spatially distinct from nucleoli. Nucleoli were labeled with Nucleolar-ID (green), and a Pepper-embedded nanostar (P-cNS3; red) was imaged in live HEK293T cells following a 30-min incubation with 5 μM HBC620 and 5 mM Mg^2+^. Dashed lines mark the nucleus of interest. Right panel shows the fluorescence intensity line-scan profiles. Scale bar: 10 μm. **b**, Confocal imaging of HEK293T cells co-expressing BFP-nucleolin with circularized Broccoli-embedded nanostar (cNS3). Cells were incubated with 40 μM DFHBI-1T and 5 mM Mg^2+^ for 30 min. Right panel shows line-scan profiles of cNS3 (green) and BFP (blue) signals. Scale bar: 10 μm. **c**, NS condensates localize adjacent to nuclear speckles. Nuclear speckles were detected by SC35 immunofluorescence (magenta), and cNS3 was visualized with 40 μM DFHBI-1T (green). Dashed lines mark the nucleus of interest. Right panels show line-scan profiles. Scale bar: 10 μm. **d**, NS condensates colocalize with paraspeckles. Paraspeckles were labeled with NONO immuno-fluorescence (magenta), and cNS3 was imaged with 40 μM DFHBI-1T (green). Line-scan profiles are shown on the right. Scale bar: 10 μm. Additional imaging demonstrates that cNS3 condensates can form even in the absence of NONO enrichment, indicating that condensate formation does not require paraspeckle assembly. **e**, Confocal imaging of HEK293T cells expressing cY2. Paraspeckles were detected with NONO (magenta), and cY2 condensates were imaged with 40 μM DFHBI-1T (green). Right panel shows fluorescence line-scan profiles. These results indicate that cY2 condensates colocalize with paraspeckles and can be capsulated by NONO-positive structure. Scale bar: 10 μm.

We then evaluated whether NS associates with nuclear speckles, which play key roles in mRNA processing and RNA metabolism^26,31–34^. cNS3 condensates did not colocalize with SC35, a nuclear-speckle marker, suggesting that NS does not interact with nuclear speckles (Fig. 4c). However, many cNS3 condensates were positioned adjacent to nuclear speckles, a localization pattern reminiscent of paraspeckles, another RNA-rich nuclear compartment involved in RNA storage and gene regulation^27,35–38^.

Given this spatial pattern, we further examined whether NS colocalizes with paraspeckles. Using NONO (p54nrb), a core paraspeckle protein, we found that cNS3 condensates largely overlapped with NONO-positive foci (Fig. 4d). Some cNS3 condensates, however, showed little or no NONO enrichment, suggesting that their association with paraspeckles may not be specific. cY2 condensates also colocalized with NONO, and in some cases NONO-positive vesicles appeared to encapsulate the condensates (Fig. 4e), further indicating a potentially indirect interaction. Because core paraspeckle components—including NONO, SFPQ, and PSPC1—are RBPs that bind RNA with complexed structure^37,38^, it is likely that dsRNA stems within NS provide multiple binding sites that recruit RBPs and promote ribonucleoprotein (RNP) condensate assembly. This mechanism would not require kissing-loop interactions and resembles how the long non-coding RNA NEAT1 scaffolds paraspeckle formation^39,40^.

### Engineering orthogonal RNA condensates through scaffold and sequence design

One intriguing advantage of RNA condensate scaffolds is their ability to form distinct, immiscible condensates, enabling precise spatiotemporal control of molecular organization^9,10,16^. To evaluate this orthogonality *in vitro*, we compared several RNA scaffold combinations. Mixing 2dB-tagged CAG repeats (20×, 31×, or 47×) with DNB-tagged 29×G_4_C_2_ produced fully merged condensates, indicating a lack of orthogonality (Fig. 5a, Supplementary Fig. 6a,b). In contrast, Broc-NS mixed with DNB-29×G_4_C_2_ formed immiscible condensates, with Broc-NS remaining peripheral (Fig. 5b). Applying a melt-and-hold protocol further enhanced this immiscibility, resulting in DNB-29×G_4_C_2_ clustering around Broc-NS condensates (Supplementary Fig. 6c).

**Fig. 5.**
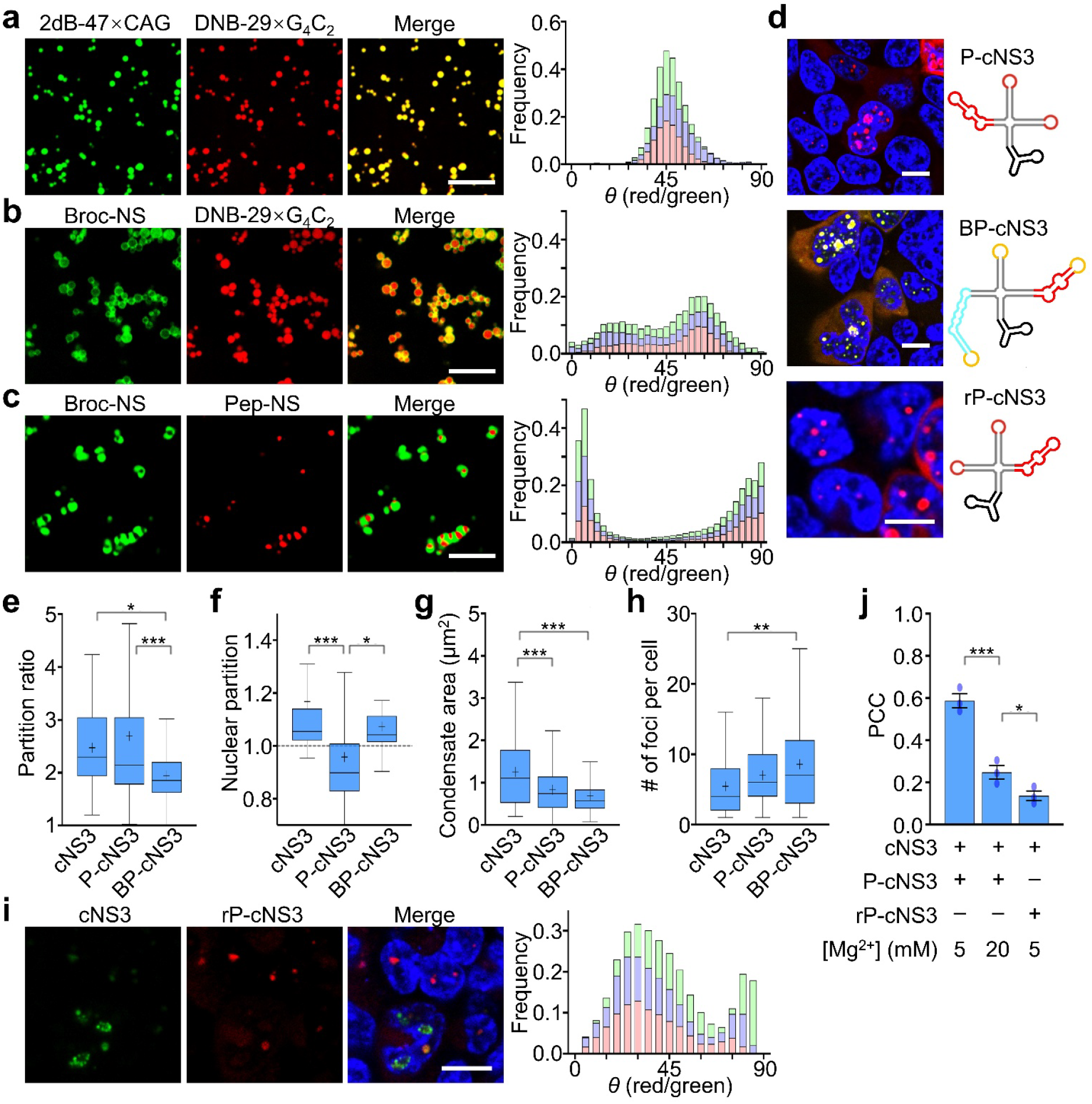
Engineering and improving orthogonal RNA nanostar condensates in cells. **a**–**c**, *In vitro* condensate formation and miscibility of RNA scaffold pairs. RNA scaffolds included 2d×Broccoli-fused 47×CAG, DNB-fused 29×G_4_C_2_, and Pepper-or Broccoli-modified nanostars (Pep-NS, Broc-NS). Equimolar scaffold pairs (2 μM each) were mixed in 20 mM MgCl_2_ with appropriate dyes (80 μM DFHBI-1T, 2 μM HBC620, or 0.5 μM TMR-DN). Fluorescence images were acquired after annealing (95 °C for 2 min, then cooled to room temperature at –0.5 °C/min; panel **a, b**), or after a “melt-and-hold” protocol (70 °C for 10 min, rapid cooling to 37 °C, then incubated for 6 h; panel **c**). Condensate orthogonality was quantified using coordinate-angle mapping from the pixel-intensity ratios of red (DNB or Pepper) and green (Broccoli) channels. Overlapping signals show a single peak at ∼45°, whereas well-separated scaffolds yield two distinct peaks at ∼0° and ∼90°. Histograms represent three independent replicates. **d**, Schematics of circular nanostars (cNS) with distinct stems and kissing-loop sequences, producing Pepper-tagged (P-cNS3) or dual-tagged Broccoli/Pepper (BP-cNS3) scaffolds. A reversed-stem nanostar, rP-cNS3, is created by inverting the cNS3 stem sequence while retaining Pepper and distinct kissing loops. This architecture is intended to reduce stem–stem interactions and enhance orthogonality with cNS3. Corresponding confocal images in HEK293T cells were acquired after incubation with 40 μM DFHBI-1T, 5 μM HBC620, and 5 mM Mg^2+^. Hoechst 33342 marks nuclei. Scale bar: 10 μm. **e**–**h**, Partition ratio, condensate area, condensate number per cell, and nuclear partition for cNS3, P-cNS3 and BP-cNS3, displayed as Tukey box-and-whisker plots. Data represents 65, 71, and 75 cells, respectively, from ≥3 independent replicates. Statistical significance was assessed by one-way ANOVA with Tukey’s post-hoc test (\*\*\**p* ≤ 0.001; \*\**p* ≤ 0.01; \**p* ≤ 0.05). Nonsignificant comparisons (*p* > 0.05) are omitted. **i**, Confocal imaging of HEK293T cells expressing rP-cNS3 (red) and cNS3 (green). The two scaffolds form distinct condensates with improved orthogonality. Imaging conditions were as in **d**. Scale bar: 10 μm. Coordinate-angle analysis of condensates shows two distinct peaks corresponding to improved orthogonality. Histograms represent three independent replicates. **j**, Pearson correlation coefficients (PCCs) quantifying orthogonality improvement. PCCs were computed from three independent replicates for cells co-transfected with cNS3 and either P-cNS3 or rP-cNS3. Values are shown as Fisher Z-transformed mean and standard deviation (SD). Statistical comparisons were performed using two-tailed Student’s *t*-test (\*\*\**p* ≤ 0.001; \**p* ≤ 0.05).

To test whether NS scaffolds could achieve stronger orthogonality, we engineered a Pepper-modified NS (Pep-NS) with distinct stem sequences and orthogonal kissing loops (Supplementary Fig. 1). After melt-and-hold incubation, Broc-NS and Pep-NS formed clearly separated condensates (Fig. 5c). Quantitative analysis using coordinate-angle mapping and Pearson correlation coefficients (PCC) showed a single *θ*-peak near 45° and PCC > 0.6 for CAG/G_4_C_2_ mixtures (Fig. 5a, Supplementary Fig. 6a,b,e), consistent with merged condensates. Broc-NS mixed with DNB-29×G_4_C_2_ showed moderate orthogonality (Fig. 5b), whereas Broc-NS and Pep-NS displayed distinct *θ*-peaks near 0° and 90° and strong negative correlation(PCC ≈ –1), indicating robust orthogonality (Fig. 5c, Supplementary Fig. 6d,e). Overall, these results suggest that RNA repeat-based scaffolds readily form merged condensates, likely due to promiscuous multivalent interactions, while NS scaffolds can generate immiscible, orthogonal condensates driven by their specific kissing-loop interactions.

Next, we examined whether NS orthogonality could be maintained in cells. To express multiple NS variants, we converted Pep-NS into P-cNS3 using the circular F30 holder and engineered BP-cNS3, which contains a distinct stem sequence and orthogonal kissing loops incorporating both Broccoli and Pepper (Fig. 5d). In HEK293T cells, both P-cNS3 and BP-cNS3 formed condensates detectable through their respective aptamer signals (Fig. 5d). Compared to cNS3, p-cNS3 exhibited similar partition ratio and condensate number but reduced nuclear enrichment and smaller condensate size. BP-cNS3 showed lower partition ratio and smaller condensates but increased condensate number (Fig. 5e–h).

To assess cellular orthogonality, we co-expressed cNS3 and P-cNS3. The two scaffolds formed merged nuclear condensates, displaying both Broccoli and Pepper signals with a single *θ*-peak and PCC of ∼0.6 (Supplementary Fig. 7), indicating loss of orthogonality. This merging likely results from nonspecific binding of nuclear RBPs to dsRNA, which draws different NS scaffolds into shared RNP assemblies. We reasoned that strengthening specific RNA–RNA interactions could counteract these nonspecific associations. Indeed, supplementing the imaging medium with 20mM Mg^2+^—which stabilizes RNA folding and base pairing^41,42^—partially restored orthogonality: cells displayed some Broccoli-only and Pepper-only condensates, and the PCC decreased to ∼0.2 (Supplementary Fig. 7).

We also sought to further improve NS orthogonality in cells. The merged condensates observed in cells—and the immiscible but contacting condensates *in vitro*—may arise from stem–stem interactions between different NS scaffolds, where segments of one stem can hybridize with another instead of self-assembling. Reducing these unintended interactions could decrease co-assembly and enhance orthogonality. To this end, we engineered a reversed cNS3 variant, replacing Broccoli with Pepper and incorporating the P-cNS3 kissing loop, generating rP-cNS3 (Fig. 5d). rP-cNS3 carries a reversed stem sequence relative to cNS3, thereby minimizing stem complementarity. When expressed alone, rP-cNS3 formed condensates in cells (Fig. 5d). When co-expressed with cNS3, the two scaffolds produced more isolated condensates (Fig. 5i), exhibiting two distinct *θ*-distributing peaks, and showed a PCC below 0.2 (Fig. 5j), indicating moderate orthogonality. Overall, these results demonstrate that careful sequence design can improve cellular NS orthogonality by enhancing specific homotypic interactions while reducing nonspecific cross-scaffold interactions.

### NS condensates enable targeted recruitment and functional control of protein and RNA clients

To enable programmable functionality, synthetic condensates must selectively recruit defined client molecules in cells. We therefore tested whether NS scaffolds can recruit client proteins into cellular condensates. P-cNS3 was modified to incorporate a protein-binding RNA hairpin on the second F30 arm, enabling interaction with proteins containing the corresponding RNA-binding domain (Fig. 6a). As a model system, we generated P-cNS3-MS2 by inserting an MS2 RNA hairpin to recruit an mNeonGreen-MS2 coat protein fusion (mNeonGreen-MCP), which contains a nuclear localization signal (NLS)^43^. When P-cNS3-MS2 and mNeonGreen-MCP were co-expressed in HEK293T cells, mNeonGreen-MCP formed foci that colocalized with P-cNS3-MS2 condensates (Fig. 6b). In contrast, mNeonGreen-MCP remained diffuse when co-expressed with P-cNS3 or expressed alone (Fig. 6b and Supplementary Fig. 8a). Although overall cellular fluorescence levels were similar (Fig. 6c), mNeonGreen-MCP exhibited a partition ratio of ∼5 when enriched within P-cNS3-MS2 condensates (Fig. 6d), demonstrating that NS scaffolds can selectively recruit client proteins through engineered RNA–protein interactions.

**Fig. 6.**
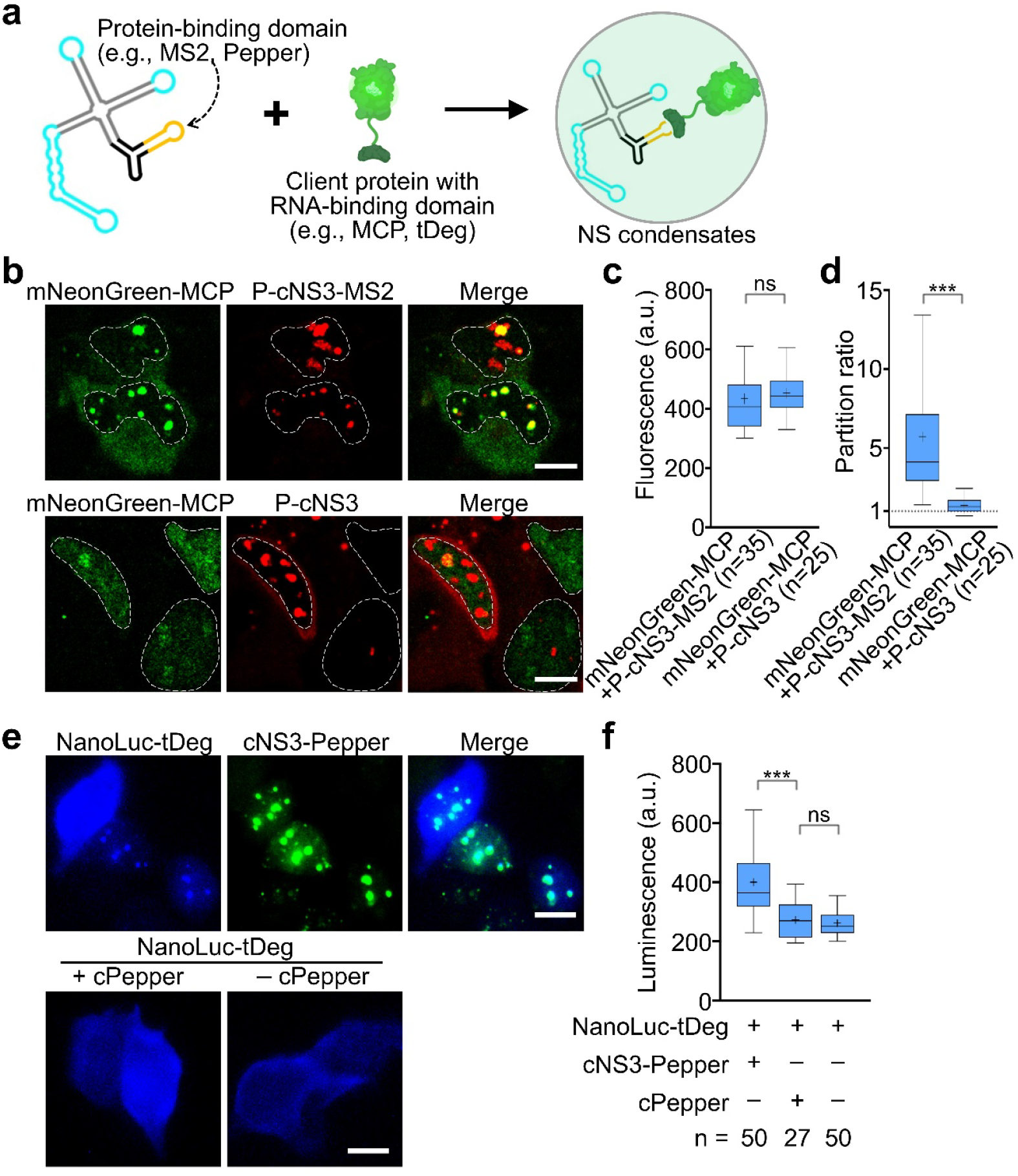
Recruitment of cellular target proteins to RNA condensates. **a**, Schematic of the modified circular nanostar (cNS3) scaffold used to recruit target proteins. The nanostar contains double-stranded stems, a fluorogenic RNA module and kissing loops, all linked to a circularized F30 three-way junction carrying a protein-binding element (MS2 hairpin or TAR Variant-2 aptamer). Client proteins contain the corresponding binding partner (MCP or tDeg). **b**, Recruitment of mNeonGreen to RNA condensates. The Pepper-embedded nanostar with an MS2 hairpin (P-cNS3-MS2) recruits mNeonGreen-MCP (upper panel). In the control (lower panel), mNeonGreen-MCP does not associate with P-cNS3. Live-cell images were acquired after a 30-min incubation with 5 μM HBC620 and 5 mM Mg^2+^. Dashed lines indicate nuclei. Scale bar: 10 μm. **c, d**, Quantification of cellular mNeonGreen fluorescence and partition ration from **b**. Data represent n cells from ≥3 independent replicates. Shown are two-tailed Student’s *t*-test (\*\*\**p* ≤ 0.001; ns, *p* > 0.05). **e**, Recruitment of NanoLuc to RNA condensates. The Broccoli-embedded nanostar containing the TAR aptamer (cNS3-Pepper) recruits NanoLuc-tDeg (upper panel). Controls include NanoLuc-tDeg expressed with or without circular TAR (cPepper). The tDeg tag promotes degradation, whereas TAR binding stabilizes the protein. Live-cell images were obtained after incubation with 40 μM DFHBI-1T, 5 mM Mg^2+^ and appropriate substrate. Scale bar: 10 μm. **f**, Cellular luminescence measurement from **e**. Background degradation can reduce luminescence differences between conditions; however, condensate recruitment increases luminescence by ∼60%. Data represent n cells from ≥3 independent replicates. Shown are two-tailed Student’s *t*-test (\*\*\**p* ≤ 0.001; ns, *p* > 0.05).

**Fig. 7.**
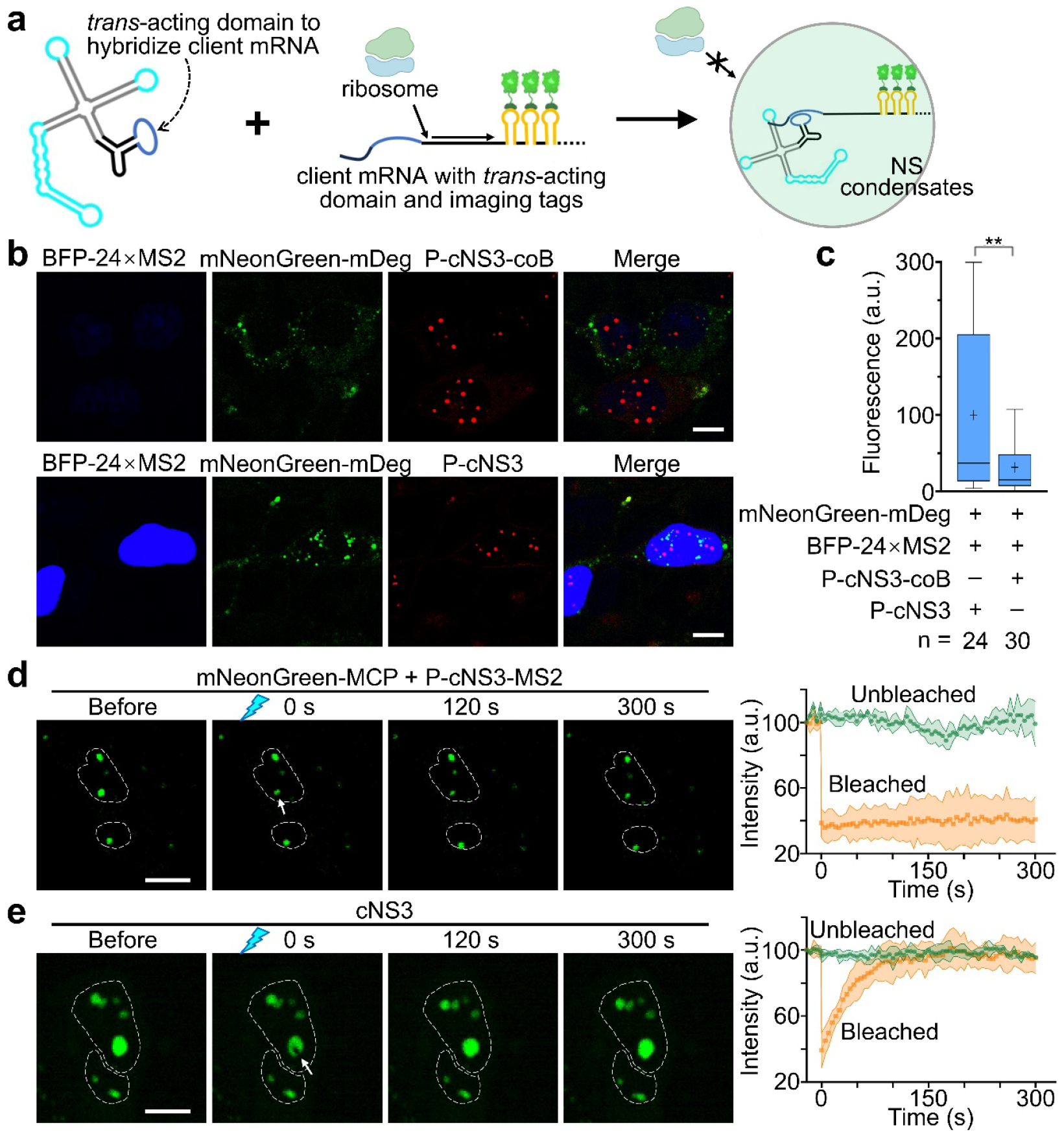
Target RNA recruitment to cellular condensates and fluorescence recovery after photobleaching (FRAP). **a**, Schematic of the modified circular nanostar scaffold containing a *trans*-acting sequence that hybridizes with and recruits target RNA into condensates. The client mRNA encoding blue fluorescent protein (BFP) carries a complementary hybridization sequence in its 5’-UTR and 24 MS2 hairpins in its 3′-UTR for visualization with mNeonGreen-mDeg. As with tDeg, the mDeg tag is degradation-prone but stabilized upon MS2 binding. Recruitment into condensates may reduce BFP translation. **b**, Reduced BFP levels upon client mRNA recruitment. The Pepper-embedded nanostar containing the *trans*-acting sequence (P-cNS3-coB) recruits BFP-24×MS2 mRNA (upper panel). mNeonGreen-mDeg visualizes the client BFP-24×MS2 RNA. In the control (lower panel), P-cNS3 does not recruit the client RNA. Live-cell images were acquired after a 30-min incubation with 5 μM HBC620 and 5 mM Mg^2+^. Scale bar: 10 μm. **c**, Quantification of relative BFP fluorescence from **b**, showing reduced BFP levels upon recruitment. Data represent n cells from ≥3 independent replicates. Shown are two-tailed Student’s *t*-test (\*\**p* ≤ 0.01). **d**, FRAP analysis showing minimal recovery of client proteins. P-cNS3-MS2 condensates recruiting mNeonGreen-MCP were photobleached and monitored for fluorescence recovery following a 30-min incubation with 5 μM HBC620 and 5 mM Mg^2+^. Dashed lines mark nuclei; white arrows indicate photobleaching regions. Scale bar: 10 μm. Right panels show FRAP trajectories (bleached condensates: orange; unbleached condensates: green) as mean ± SD from three independent replicates. **e**, FRAP analysis showing a rapid recovery of small molecules. cNS3 condensates labeled with DFHBI-1T were photobleached after a 30-min incubation with 40 μM DFHBI-1T and 5 mM Mg^2+^. Scale bar, 10 μm. Right panels show FRAP trajectories (bleached condensates: orange; unbleached condensates: green) as mean ± SD from three independent replicates.

Next, we examined how condensate recruitment influences client protein function. We engineered cNS3-Pepper, which incorporates the TAR Variant-2 “Pepper” aptamer to bind and stabilize a NanoLuc fusion protein carrying a C-terminal tDeg degron (NanoLuc-tDeg). The tDeg tag promotes protein degradation, whereas Pepper binding inhibits degradation and stabilizes the fused protein^44^. Note that this Pepper aptamer is distinct from the fluorogenic RNA aptamer described earlier. When cNS3-Pepper and NanoLuc-tDeg were co-expressed in HEK293T cells, NanoLuc-tDeg formed luminescent foci that colocalized with cNS3-Pepper condensates (Fig. 6e). Notably, compared with cells co-expressing NanoLuc-tDeg and circular Pepper (cPepper), recruitment into cNS3-Pepper condensates increased cellular luminescence by ∼60% (Fig. 6f). This enhancement may result from condensate-mediated sequestration of NanoLuc-tDeg away from proteasomal degradation or from local enrichment that enhances its enzymatic activity.

We next asked whether recruitment into condensates also affects the function of client RNAs. To text this, we modified P-cNS3 to include a *trans*-acting domain that enables client RNA binding through base pairing (Fig. 7a). As a model system, we used an exogenously expressed blue fluorescent protein mRNA (BFP-24×MS2), which contains a base-pairing domain in its 5’ UTR for condensate recruitment and 24 MS2 hairpins in its 3′ UTR for visualization. The encoded BFP also carries an NLS. The corresponding NS scaffold designed to recruit this mRNA was named P-cNS3-coB. For visualization, we used mNeonGreen-mDeg, a destabilized MCP fusion that can be stabilized by MS2 binding^45^.

When BFP-24×MS2 and mNeonGreen-mDeg were co-expressed, nuclear BFP signals and cytoplasmic mNeonGreen-mDeg-labeled mRNA foci were observed (Supplementary Fig. 8b). Co-expression of BFP-24×MS2, mNeonGreen-mDeg, and P-cNS3 produced mRNA foci that remained separate from p-cNS3 condensates (Fig. 7b). Notably, co-expression of BFP-24×MS2, mNeonGreen-mDeg, and P-cNS3-coB did not result in detectable recruitment of mNeonGreen-mDeg into condensates (Fig. 7b), likely because sequestration within condensates blocked MS2–mDeg interaction. However, BFP-24×MS2 expression decreased approximately fourfold when co-expressed with P-cNS3-coB (Fig. 7c), indicating that condensate recruitment sequesters the client mRNA away from the translation machinery and other RNA–protein interactions. Together, these findings show that the condensate environment can substantially alter the behavior and function of client RNAs.

Using the mNeonGreen-MCP/P-cNS3-MS2 system, we next evaluated the dynamics of NS condensates in cells. Fluorescence recovery after photobleaching (FRAP) of mNeonGreen-MCP-labeled P-cNS3-MS2 condensates showed no recovery over 300 s, indicating low mobility of client proteins within these RNA condensates (Fig. 7d). In contrast, FRAP of the embedded fluorogenic RNA aptamer and dye in cNS3 condensates showed rapid recovery, reaching more than 80% of the original fluorescence within 100 s, suggesting fast exchange of small-molecule clients (Fig. 7e). Similar rapid dye recovery was observed for other NS scaffolds, including cNS2, cNS0, cY2 and cY1, with ∼80% recovery within 200 s (Supplementary Fig. 9).

FRAP performed *in vitro* on linear NS3 and Y2 revealed even faster recovery—70%–80% within 5 s (Supplementary Fig. 10)—although the bleached center did not recover over 300 s, leaving ∼20% immobile fraction (Supplementary Fig. 9). This suggests the presence of a compact core region inaccessible to small molecules. Overall, these data indicate that cellular NS condensates form a crowded environment in which macromolecules such as proteins exhibit limited mobility, whereas small molecules remain freely exchangeable.

### Small-molecule-responsive RNA scaffolds enable switchable condensate formation

To enhance the programmability of RNA condensates, we sought to develop a switchable, small-molecule-responsive RNA scaffold. We incorporated an allosteric RNA switch into the NS architecture to enable acyclovir-dependent conformational changes that toggle condensate formation^46^. Conceptually, disrupting the dsRNA stem and kissing loop prevents condensation, whereas restoring them re-establishes assembly. To test this design, we engineered cNS3-Sw and cY2-Sw by integrating the acyclovir-responsive RNA nanodevice “RN4”^46^, which modulates the folding of one kissing loop and its connecting arm (Fig. 8a, b). Linear cNS3-Sw and cY2-Sw tested *in vitro* showed acyclovir-induced condensation, with ∼50% increases in partition ratio following melt-and-hold incubation (Supplementary Fig. 11a–c). cNS3-Sw also exhibited a ∼threefold increase in condensate size, whereas cY2-Sw showed minimal size change (Supplementary Fig. 11d). These results demonstrate that integrating an allosteric switch into NS scaffolds enables chemically inducible RNA condensation *in vitro*.

**Fig. 8.**
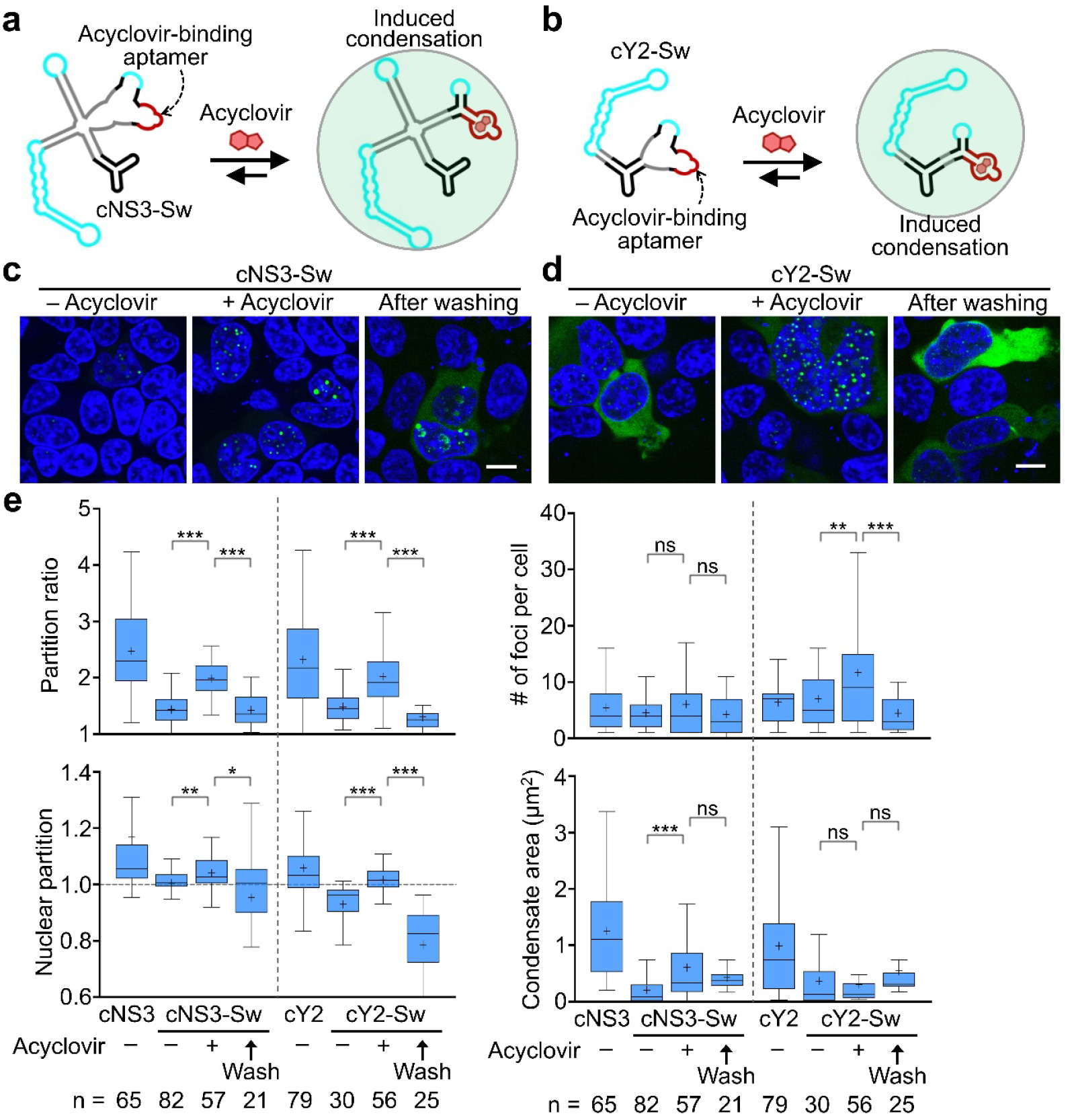
Acyclovir-induced switchable RNA condensate formation. **a, b**, Schematics of allosteric RNA switch-modified circular nanostar scaffold (cNS3-Sw and cY2-Sw). The acyclovir-responsive switch disrupts one stem and kissing loop, preventing condensate formation. Acyclovir binding restores proper folding, enabling condensate assembly. **c, d**, Acyclovir-induced assembly and disassembly of RNA condensates in HEK293T cells. Cells expressing cNS3-Sw or cY2-Sw were treated with vehicle (0.1% v/v DMSO) or 100 μM acyclovir for 4 h before imaging. To remove acyclovir, the medium was replaced with fresh medium, and cells were further incubated for an additional 2 h. Live-cell images were obtained after a 30-min incubation with 40 μM DFHBI-1T and 5 mM Mg^2+^. Hoechst 33342 marks nuclei. Scale bar: 10 μm. **e**, Quantification of partition ratio, condensate number per cell, nuclear partition, and condensate area from **c, d**, displayed as Tukey box-and-whisker plots. Boxes represent the 25th, 50th, and 75th percentiles; whiskers extend to 1.5× the interquartile range; means are shown as “+”. Data represents n cells from ≥3 independent replicates. Statistical significance was assessed using two-tailed Student’s *t*-tests (\*\*\**p* ≤ 0.001; \*\**p* ≤ 0.01; \**p* ≤ 0.05; ns, *p* > 0.05).

We next tested whether these switchable scaffolds function in living cells. In HEK293T cells, both cNS3-Sw and cY2-Sw showed acyclovir-induced condensate formation, with significant increases in partition ratio and nuclear enrichment (Fig. 8c–e). For cNS3-Sw, acyclovir primarily increased condensate size, whereas cY2-Sw showed an increase in condensate number. Importantly, the induced condensates were reversible: washing with fresh medium restored condensate properties to pre-acyclovir levels (Fig. 8c–e). Time-lapse imaging over 8 hours captured the assembly and disassembly dynamics (Supplementary Video 1, 2). Together, these findings demonstrate that NS scaffolds can be engineered to support small-molecule-controlled, reversible RNA condensate formation in living cells.

## DISCUSSION

This study establishes a genetically encoded, programmable platform for RNA-driven synthetic condensates in mammalian cells. Although RNA is recognized as a versatile scaffold^4^, robust and dynamically controllable RNA-based condensate systems have been limited. Through systematic evaluation of repeat-based and *de novo* scaffolds, we developed RNA nanostar variants (such as cNS3 and cY2) that consistently induce nuclear condensate formation. A key and unexpected finding is that, within cells, condensate assembly is driven primarily by double-stranded RNA (dsRNA) stem engaging endogenous nuclear RNA-binding proteins (RBPs), rather than by the kissing-loop interactions that dominate *in vitro* phase separation. This distinction reveals a fundamental divergence between simplified *in vitro* models and the complex intracellular environment, where pervasive RBP–dsRNA interactions can overshadow designed RNA–RNA contacts and limit scaffold orthogonality.

Recognizing the central role of RBP–dsRNA interactions reshape the design principles for RNA-based condensation. In mammalian cells, phase separation often relies on heterotypic interactions with abundant RBPs acting as co-scaffolds, rather than relying solely on designed RNA–RNA contacts. dsRNA stems likely engage in dynamic, multivalent interactions with nuclear RBPs containing dsRNA-binding domains, explaining both the limited orthogonality observed in cells and the challenge of achieving sequence-specific condensate behavior. By refining scaffold sequences to minimize nonspecific stem–stem pairing and strengthen specific homotypic interactions, we improved orthogonality in cells, though future work will be needed to identify dsRNA elements or scaffolds that better resist incorporation into endogenous RNP networks.

Beyond establishing scaffold performance, we demonstrated that nanostar condensates function as operational cellular compartments. Recruitment of protein and mRNA clients modulated their stability and activity. Sequestering NanoLuc increased its luminescence, whereas recruiting BFP mRNA suppressed its translation, illustrating how condensates can concentrate or isolate biomolecules to alter their interactions landscapes, stability, and activity. By swapping binding domains, this framework can be readily adapted to target diverse molecules, expanding its potential for both synthetic biology and therapeutic applications.

We further expanded the functional scope of the platform by introducing an acyclovir-responsive allosteric switch to achieve reversible, small-molecule-controlled condensation. This integration highlights the potential of allosteric RNA devices to generate stimulus-responsive biomaterials in cells, with precise spatiotemporal regulation. Although we have demonstrated controllable assembly and disassembly, additional optimization may enhance both the robustness of induction and the completeness of condensate dissolution. Replacing the RNA switch module could also broaden the range of triggers—such as light, metabolites, or other signaling molecules—thereby increasing applicability across biological contexts.

Collectively, by providing a toolkit of engineered RNA scaffolds, this work enables the construction of synthetic cellular compartments whose formation, composition, and dissolution can be genetically programmed and chemically regulated. These capabilities open avenues for fundamental studies of RNP assembly and intracellular compartmentalization, as well as for applications such as biosensing and therapeutic strategies that direct specific targets into designed condensates^6^. During the preparation of this manuscript, we became aware of a recent preprint from the Franco Lab that also reports the construction of artificial RNA nanostar condensates in mammalian cells^47^. Both works together underscore RNA’s dual capacity as both an informational polymer and a structural material, positioning synthetic RNA condensates as a versatile and programmable interface for cellular engineering. Future efforts should focus on defining the RBP interactomes associated with each scaffold and developing hybrid designs that integrate selective RNA–RNA and RNA–protein interactions to achieve orthogonal and functional condensation in cells.

## METHODS

### Materials and general methods

All chemicals were obtained from MilliporeSigma or Fisher Scientific unless otherwise noted. Commercial reagents were used as received without further purification. HBC620, 4-(2-hydroxyethyl-methylamino)-benzylidene-cyanophenylacetonitrile analog, was purchased from GlpBio (cat. GC60186); 3,5-difluoro-4-hydroxybenzylidene imidazolinone-2-oxime (DFHO) from Bio-techne (cat. 6434); and tetramethyl-rhodamine-2,4-dinitroaniline (TMR-DN) from Lumiprobe (cat. 2641).

Single-stranded DNA oligonucleotides were purchased and cartridge-purified from Integrated DNA Technologies (Coralville, IA) or the W. M. Keck Oligonucleotide Synthesis Facility (Yale University School of Medicine). Oligonucleotides were then dissolved at 100 μM concentration in syringe-filtered TE buffer (10 mM Tris–HCl, 0.1 mM EDTA, pH 7.5) and stored at –20°C. Double-stranded DNA fragments (gBlocks) were obtained from Integrated DNA Technologies and reconstituted in TE buffer following the manufacturer’s instructions.

Double-stranded DNA templates and inserts were generated by PCR amplifications in an Eppendorf Mastercycler using Q5 High-Fidelity 2× Master Mix (New England Biolabs, NEB, M0492) and purified with the Monarch PCR & DNA Cleanup Kit (NEB, T1030). Nucleic acid concentrations were measured using a NanoDrop One UV-vis spectrophotometer. PCR products were further purified by 1% (w/v) Tris-acetate-ethylenediaminetetraacetic acid (TAE) agarose gel electrophoresis followed by extraction with the Thermo Scientific Gel Extraction kit (cat. FERK0692).

Restriction digests of PCR products and DNA plasmids were performed using NEB restriction endonucleases, and DNA ligations were assembled with Gibson Assembly Master Mix (NEB, E2611). Plasmids were propagated in 5-alpha Competent *E. coli* (NEB, C2987) or Stable Competent *E. coli* (NEB, C3040) for constructs containing repeat sequences, and extracted using GeneJET Plasmid Miniprep Kit (Thermo Scientific, FERK0503). Sequence identity of DNA plasmids was confirmed via Sanger sequencing (Eurofins Genomics) or whole-plasmid sequencing (Azenta Life Sciences or Plasmidsaurus).

### *In vitro* condensate formation and imaging

RNAs were synthesized via *in vitro* transcription using a HiScribe T7 high-yield RNA synthesis kit (NEB, E2040). Following transcription, DNA templates were removed with DNase I (NEB, M0303), and RNA products were purified using G-25 columns and verified by 10% denaturing polyacrylamide gel electrophoresis (PAGE). All the purified RNA products were aliquoted and stored at –20 °C for short-term use or –80°C for long-time storage. RNA structures were designed and simulated using NUPACK and mFold online software.

*In vitro* RNA condensate formation followed a previously described protocol^13^ unless otherwise specified. Briefly, 4 μM RNA in buffer (10 mM Tris–HCl, 100 mM KCl, 20 mM MgCl_2_, pH 7.5) was heated up to 95 °C for 2 min and cooled to 37°C at a rate of –0.5 °C/min in a thermocycler. Appropriate dye (80 μM DFHBI-1T, 2 μM HBC620, 0.5 μM TMR-DN or 2 μM DFHO) was then added and incubated for 30 min at 37 °C before imaging. For RNA nanostar scaffolds, a modified “melt-and-hold”^10^ procedure was used. Briefly, 4 μM RNA in buffer (40 mM HEPES, 100 mM KCl, 500 mM NaCl, 20 mM MgCl_2_) was heated at 70 °C for 10 min, rapidly cooled to 37 °C, and incubated at 37 °C for 6, 12 or 24 hours in a thermocycler. Appropriate dyes were added 30 min before incubation at 37 °C. To induce switchable nanostar condensate formation, samples were treated with either 100 μM acyclovir (in 0.1% v/v DMSO) or a 0.1% v/v DMSO vehicle control prior to the “melt-and-hold” incubation.

### *In vitro* condensate imaging

Imaging was performed at room temperature using a Yokogawa spinning-disk confocal system on a Nikon Eclipse-TI inverted microscope using a 100×/1.45 NA oil-immersion objective. Broccoli/DFHBI-1T, Corn/DFHO, and Beetroot/DFHO were excited with a 488 nm laser (emission 500–550 nm), while Pepper/HBC620 and DNB/TMR-DN were imaged using a 561 nm laser (emission 575–625 nm). Image analysis was conducted using the General Analysis 3 (*GA3*) module in NIS-Elements AR Analysis software. Briefly, a fluorescence *Segmentation > Threshold* F_B_ + 3SD (background plus 3-fold of standard deviations on the imaging channel) and a *Threshold > Size* of 0.5 μm was set to automatically detect each condensate. All segmentations were manually verified to avoid inappropriate detection. For each identified condensate objects, the *Object Intensity > Mean, Object Area > Mean* and *Object Elongation > Mean* in *Measurement* nodes were applied to measure their corresponding mean fluorescence intensity, condensate area and aspect ratio, respectively. Mean background intensity was measured using an inverted threshold set, and the partition ratio was calculated as the mean fluorescence intensity within each condensate divided by the solution background fluorescence.

### Cell culture and transfection

HEK293T/17 (CRL-11268), HeLa (CCL-2), and MCF-7 (HTB-22) cells were purchased from American Type Culture Collection (ATCC). Cells were cultured in Dulbecco’s modified Eagle’s medium (Thermo Scientific, 11995-065) supplemented with 10% fetal bovine serum (Thermo Scientific, A5670801), 100 U/mL of penicillin, and 100 μg/mL of streptomycin (Thermo Scientific, 15140122) at 37 °C with 5% CO_2_. All cell lines were confirmed to be mycoplasma-free, and cell passage was performed using TrypLE Express (Thermo Scientific, 12604013) at ∼80% confluence.

For transfection, cells were seeded at a density of 3.2 × 10^5^ cells per 35-mm poly-D-lysine-coated glass-bottom dish (Cellvis, D35-20-1.5H) and cultured overnight. Transfections were performed the following day using FuGENE HD (Promega, E2312) according to the manufacturer’s protocol. Briefly, 2.8 µg of total DNA plasmids (1:1 mass ratio for co-transfections) were mixed in 125 μL Opti-MEM medium (Thermo Scientific, 31985062) with 6.4 μL FuGENE HD, and incubated at room temperature for 15 min. The resulting complex was then added dropwise to each dish. Cells were imaged 48 hours post-transfection.

### Vector construction

Plasmids expressing circular RNA scaffolds were generated based on a pAVU6+27-Tornado vector (Addgene, 124360), which contains a U6 promoter and two self-cleaving ribozyme sequences (TTRz and TRz) for autocatalytic RNA circularization. The Tornado DNA vector was double digested by NotI-HF (NEB, R3189) and SacII (NEB, R0157) to prepare the Gibson assembly backbone. Inserts were generated by PCR amplifications or obtained as gBlock from Integrated DNA Technologies, each designed with 25-bp overlaps at both ends for Gibson assembly. Gibson Assembly Master Mix (NEB, E2611) was used following manufacturer’s protocol. Additional plasmids used in this study included: pcDNA3.1-CMV-TagBFP-hNucleolin (Addgene 182592), pCDH-CMV-NanoLuc-tDeg, pcDNA3.1-miniCMV-4×mNeonGreen-NLS-stdMCP (mNeonGreen-MCP), pcDNA3.1-CMV-NLS-mTagBFP2-24×MS2 (BFP-24×MS2), pcDNA3.1-miniCMV-4×mNeonGreen-mDeg (mNeonGreen-mDeg), pAV-U6+27-tnd-5×Corn, and pAV-U6+27-tnd-5×Beetroot (gifts from the Jiahui Wu laboratory at UMass Amherst). All plasmid sequences are provided in the List of plasmid Sequences.

### Live-cell fluorescence and luminescence imaging of RNA condensates

Live-cell imaging was performed using a Yokogawa spinning-disk confocal system on the Nikon Eclipse-TI inverted microscope with a 40×/1.3 NA oil-immersion objective. Before imaging, the cell culture medium was replaced with phenol-red-free DMEM (Corning, MT17205CV) supplemented with 5 mM MgCl_2_ and specific fluorogenic dyes: 40 μM DFHBI-1T, 5 μM HBC620, 1 μM TMR-DN, or 40 μM DFHO. Cells were incubated at 37 °C for 30 min post-staining and subsequently imaged at room temperature. Nuclei were visualized using Hoechst 33342 (Thermo Scientific, NucBlue, R37605). Nucleoli were visualized using NUCLEOLAR-ID Green Detection Kit (Enzo Biochem, ENZ-51009). Fluorescence signals were collected using standard laser channel: Pepper/HBC620 and DNB/TMR-DN (561 nm excitation, 575–625 nm emission), Broccoli/DFHBI-1T, Corn/DFHO and mNeonGreen (488 nm excitation, 500–550 nm emission), and Hoechst 33342 and BFP (405 nm excitation, 425–475 nm emission).

To induce switchable nanostar condensates, HEK293T cells expressing cNS3-Sw or cY2-Sw were treated with either 100 μM acyclovir (in 0.1% v/v DMSO) or a 0.1% v/v DMSO vehicle control for 4 h. For acyclovir washout, the medium was replaced with phenol-red-free DMEM containing 40 μM DFHBI-1T and 5 mM MgCl_2_ followed by a 2 h incubation prior to further imaging. Time-lapse imaging of switchable nanostar condensate assembly and disassembly was performed using a Nikon Ti2-E epifluorescence microscope equipped with a Prime BSI Express sCMOS camera and a Lumencor SOLA V-NIR Light Engine. Cells were maintained at 37 °C and 5% CO_2_ using a Tokai Hit stage-top incubator. Following a 4-h treatment with 100 μM acyclovir, the medium was replaced, and imaging continued for an additional 4 h. Images were acquired every 10 min over an 8-h observation period.

Image analysis was performed using the *GA3* module in NIS-Elements AR Analysis software. Briefly, *Image Processing > Smooth* was applied to all channels before segmentation. A fluorescence intensity *Segmentation > Threshold* F_B_ + 3SD (whole dish background plus 3-fold of standard deviations) on the bule channel was set to detect each Hoechst-stained nucleus. To detect intracellular foci, the *Segmentation > Bright Spots* was applied after an intensity *Threshold* F_B_ + 3SD (cellular background plus 3-fold of standard deviations on the green or red foci channel) and a *Threshold > Size* of 0.5 μm. *Bright Spots > Contrast* and *Grow* were also set to enable the proper detection of foci. *Binary Processing > Watershed* was applied on foci channel for cellular boundary segmentation. The cell foci detection and boundary of each cell was manually verified and corrected. *Binary Processing > Make Cell* was then applied to associate foci and nucleus with individual cell. For each identified cell, the *Nucleus Intensity > Mean, Cytoplasm Intensity > Mean, Cell Intensity > Mean, Spot Intensity > Mean, Spot Area > Mean* and *Spots Count* were measured under the *Measurement > Cell* node. Partition ratio was calculated as the ratio of mean *Spot Intensity* versus the mean *Cell Intensity* of the corresponding cell. Nuclear partition was calculated as the ratio of mean *Nucleus Intensity* versus the mean *Cytoplasm Intensity*. Mean *Spot Area* and *Spots Count* generated the values for condensate area and condensate number per cell, respectively. In orthogonality measurement, pixel intensity of the segmented foci from two channels was exported via *Measurement > Pixel Values* node. Pearson correlation coefficients (PCC) were calculated using these pixel intensities. Coordinate-angle mapping was performed to visualize orthogonality^10^. In brief, a coordinate angle *θ* for each pixel was calculated as the arctangent of the pixel-intensity ratio of two foci channels. Frequency distribution histogram of angle *θ* was graphed with 5°-bin width.

For NanoLuc luminescence imaging, HEK293T cells were seeded at 3 × 10^4^ cells per well in poly-D-lysine-coated 8-well chambered coverglass plates (Cellvis, C8-1.5H-N) and cultured overnight. Transfection was performed the following day using 2.8 µg of total DNA plasmids (1:1 mass ratio for co-transfections) in each well via FuGENE HD according to the manufacturer’s protocol. These cells were imaged 24 hours after transfection. Cell culture media was changed to DPBS containing 40 μM DFHBI-1T and cultured at 37 °C for 60 min prior to imaging. Images were acquired immediately after addition of furimazine (1% v/v) using a Nikon Ti2E motorized inverted microscope equipped with a Prime BSI Express sCMOS monochrome camera via a 40×/0.95 NA air objective at 23 °C. Broccoli/DFHBI-1T fluorescence was collected using a GFP filter cube with excitation filter 470 ± 20 nm, dichroic mirror 495 nm (long pass) and emission filter 525 ± 25 nm. Luminescence was collected using a 460 ± 25 nm emission filter with a 10-s exposure time.

### Cell viability assay

HEK293T cell proliferation was assessed using the CyQUANT XTT Cell Viability Assay (Invitrogen, X12223). Cells were seeded in 96-well plates (Thermo Scientific, 165305) at 1 × 10^4^ cells per well in 100 μL of complete medium and allowed to adhere overnight. The cells were then transfected with plasmids encoding various RNA nanostar scaffolds (cNS3, cNS2, cNS0, cY2, cY1, cuNS3, cuNS0, csNS3, csNS0, or c0) using FuGENE HD according to the manufacturer’s instructions. Control groups included untransfected cells (no treatment) and mock-transfected cells (FuGENE HD reagent without plasmid). Following a 24-hour incubation at 37°C, XTT working solution was prepared by mixing 6 mL of XTT reagent with 1 mL of electron coupling reagent; 70 μL of this mixture was added to each well. After a further 4-hour incubation at 37°C, the absorbance was measured at 450 nm and 660 nm (background) using a BioTek Synergy 2 microplate reader. Relative cell viability was determined by normalizing the background-corrected absorbance of the experimental groups to that of the untransfected controls.

### Immunofluorescence

Cells were seeded and transfected in 8-well chambered plates (Cellvis, C8-1.5H-N). After rinsing twice with PBS (Thermo Fisher, 10010023), cells were fixed with 4% paraformaldehyde (Thermo Fisher, AAJ61899AP) in PBS for 15 min at room temperature, followed by three 5-min washes with PBS. Permeabilization was performed using 0.1% Triton X-100 (Sigma Aldrich, 648462) in PBS for 5 min at room temperature, followed by three 5-min PBS washes. Nonspecific binding was blocked for 30 min at room temperature using a blocking buffer containing 5% BSA (Sigma Aldrich, 05470), 5% goat serum with 0.05% sodium azide (Thermo Fisher, 50197Z), and 0.1% Tween 20 (Sigma Aldrich, P1379) in PBS. For immunofluorescence, cells were incubated with primary antibodies—mouse anti-SC35 (Abcam, ab11826) or mouse anti-p54nrb (BD Biosciences, BDB611278)—at 5 μg/mL in dilution buffer (PBS containing 1% BSA and 0.05% Tween 20) for 60 min at room temperature. After three 5-min PBS washes, secondary antibodies (Abcam, ab150115) were applied at 10 μg/mL in dilution buffer for 60 min at room temperature. Following a final set of three 5-min washes, cells were incubated in PBS supplemented with 40 μM DFHBI-1T and NucBlue reagent for 30 min prior to imaging.

### Fluorescence recovery after photobleaching (FRAP)

FRAP experiments were performed using a Yokogawa spinning-disk confocal system on the Nikon Eclipse-TI inverted microscope. Samples were imaged using a 100×/1.45 NA oil-immersion objective for *in vitro* experiments and a 40×/1.3 NA objective for live-cell imaging. Photobleaching was executed with a 488 nm laser for 1.0 s over a ∼1 μm diameter region. Following a single pre-bleach frame, images were acquired every 5 s for 6 min with 2×2 binning. Mean fluorescence intensities of the bleached and nearby unbleached (control) condensates were quantified in ImageJ and normalized to their respective pre-bleach values. Data are presented as mean ± standard deviation values from three independent replicates.

### Statistical analysis

Data visualization and statistical analyses were performed using GraphPad Prism and Microsoft Excel. Sample sizes (n) for each experiment are indicated in the figures or corresponding legends. All measurements were derived from at least three independent replicates unless otherwise specified. Statistical significance was determined using unpaired, two-tailed Student’s *t*-tests for pairwise comparisons or one-way analysis of variance (ANOVA) followed by Tukey’s post-hoc test for multiple comparisons.

## Acknowledgements

The authors gratefully acknowledge the support from NSF 2435059, Chan Zuckerberg Initiative Dynamic Imaging program 2023-321170, and Camille Dreyfus Teacher-Scholar Award to M.Y. We would also like to thank the Society for Laboratory Automation and Screening (SLAS) for providing the graduate education fellowship grant to L.M, and want to thank NIH Traineeship T32GM139789 to R.Z. The authors also thank Prof. Jiahui Wu and other members of the You Lab for useful discussion and valuable comments.

## Author contributions

Conceptualization: Z.X. and M.Y. Methodology: Z.X., R.Z. and M.Y. Visualization: Z.X., O.M. and M.Y. Data curation: Z.X., O.M., L.M. and M.Y. Investigation: Z.X., O.M., L.M., Y.G. and C.Y. Formal analysis: Z.X., O.M. and M.Y. Funding acquisition & Resources: M.Y. Supervision: M.Y. Writing-original draft: Z.X. and M.Y. Writing-review & editing: Z.X. and M.Y.

## Additional information

Supplementary information and source data that support the findings of this study are available in additional documents and will also be openly available online.

