## Supplementary Information for "Programmable and Switchable RNA Scaffolds for Synthetic Condensate Engineering in Mammalian Cells"

### SUPPLEMENTARY TABLE

**Table S1.** List of RNA sequences used in this study. Blue nucleotides indicate the sequences of F30 scaffold and its variants. Turquoise highlights the kissing-loop binding regions, while the grey highlights the inert kissing-loop regions. Red nucleotides are ribozyme cutting sites for RNA cyclization in the Tornado vector. Underlined nucleotides show the *trans*-acting hybridization domain. Orange nucleotides indicate the sequences of acyclovir-responsive switch.

| Name | RNA sequences (5’-3’) |
|---|---|
| 2d× Broccoli (2dB) | GACUUGCCAUGUGUAUGUGGGAGACGGUCGGGUCCAUCUGAGACGGUCGGGUCCAGA<br>UAUUCGUAUCUGUCGAGUAGAGUGUGGGCUCAGAUGUCGAGUAGAGUGUGGGCUCCC<br>ACAUACUCUGAUGAUCCAGACGGUCGGGUCCAUCUGAGACGGUCGGGUCCAGAUAUU<br>CGUAUCUGUCGAGUAGAGUGUGGGCUCAGAUGUCGAGUAGAGUGUGGGCUGGAUCAU<br>UCAUGGCAA |
| 2dB-20×CAG | GACUUGCCAUGUGUAUGUGGGAGACGGUCGGGUCCAUCUGAGACGGUCGGGUCCAGA<br>UAUUCGUAUCUGUCGAGUAGAGUGUGGGCUCAGAUGUCGAGUAGAGUGUGGGCUCCC<br>ACAUACUCUGAUGAUCCAGACGGUCGGGUCCAUCUGAGACGGUCGGGUCCAGAUAUU<br>CGUAUCUGUCGAGUAGAGUGUGGGCUCAGAUGUCGAGUAGAGUGUGGGCUGGAUCAU<br>UCAUGGCAAUCUAGAGAAUUCGGUCUCACAGCAGCAGCAGCAGCAGCAGCAGCAGCAG<br>CAGCAGCAGCAGCAGCAGCAGCAGCAGCAGUUUUUUCCGCGG |
| 2dB-31×CAG | GACUUGCCAUGUGUAUGUGGGAGACGGUCGGGUCCAUCUGAGACGGUCGGGUCCAGA<br>UAUUCGUAUCUGUCGAGUAGAGUGUGGGCUCAGAUGUCGAGUAGAGUGUGGGCUCCC<br>ACAUACUCUGAUGAUCCAGACGGUCGGGUCCAUCUGAGACGGUCGGGUCCAGAUAUU<br>CGUAUCUGUCGAGUAGAGUGUGGGCUCAGAUGUCGAGUAGAGUGUGGGCUGGAUCAU<br>UCAUGGCAAUCUAGAGAAUUCGGUCUCACAGCAGCAGCAGCAGCAGCAGCAGCAGCAG<br>CAGCAGCAGCAGCAGCAGCAGCAGCAGCAGCAGCAGCAGCAGCAGCAGCAGCAGCAGC<br>AGCAGUUUUUUCCGCGG |
| 2dB-47×CAG | GACUUGCCAUGUGUAUGUGGGAGACGGUCGGGUCCAUCUGAGACGGUCGGGUCCAGA<br>UAUUCGUAUCUGUCGAGUAGAGUGUGGGCUCAGAUGUCGAGUAGAGUGUGGGCUCCC<br>ACAUACUCUGAUGAUCCAGACGGUCGGGUCCAUCUGAGACGGUCGGGUCCAGAUAUU<br>CGUAUCUGUCGAGUAGAGUGUGGGCUCAGAUGUCGAGUAGAGUGUGGGCUGGAUCAU<br>UCAUGGCAAUCUAGAGAAUUCGGUCUCACAGCAGCAGCAGCAGCAGCAGCAGCAGCAG<br>CAGCAGCAGCAGCAGCAGCAGCAGCAGCAGCAGCAGCAGCAGCAGCAGCAGCAGCAGC<br>AGCAGCAGCAGCAGCAGCAGCAGCAGCAGCAGCAGCAGCAGCAGCAGCAGCAGUUUUU<br>UUCCGCGG |
| Broc-47×CUG | GCGGCCGCUUGCCAUGUGUAUCGGGGAGACGGUCGGGUCCAGAUAUUCGUAUCUGUC<br>GAGUAGAGUGUGGGCUCCCCGAUACUCUGAUGAUCCUCUAGAGAAUUCCUGCUGCUG<br>CUGCUGCUGCUGCUGCUGCUGCUGCUGCUGCUGCUGCUGCUGCUGCUGCUGCUGCU<br>GCUGCUGCUGCUGCUGCUGCUGCUGCUGCUGCUGCUGCUGCUGCUGCUGCUGCUGC<br>UGCUGCUGCUGCUGCUGCUGGCUAGCAUAAGGAUCAUUCAUGGCAA |
| Pep-47×CAG | GCGGCCGCUACCGUAGGCUCGCCGUGUGUGUGUAGGAGGGAUCCCCAAUCGUGGCG<br>UGUCGGCCUGCUUCGGCAGGCACUGGCGCCGGGAUCCCUCCUACACACUCUGACGAU<br>CCUCAUUCAUGGCAAUCUAGAGAAUUCGGUCUCACAGCAGCAGCAGCAGCAGCAGCAG<br>CAGCAGCAGCAGCAGCAGCAGCAGCAGCAGCAGCAGCAGCAGCAGCAGCAGCAGCAGC<br>AGCAGCAGCAGCAGCAGCAGCAGCAGCAGCAGCAGCAGCAGCAGCAGCAGCAGCAGCA<br>GUUUUUUCCGCGGCUAGCAUAACCCCUUGGGAUCGUUCACGGCGAGCCUACGGUA |
| DNB-29×$G_4C_2$ | GGUUGCCAUGUGUAUCGGGUGCCUUAUUCCGGACGCCGGGCCCGAAUGCUGCUACGG<br>CAGUCGAAGACAACAUCGCGCCCUUCGGAGGCACCCGAUACUCUGAUGAUCCUUGAAU<br>UCGCUAGCUAACUAGCUAACUAGGCUCUUCCGGCCGGGGCCGGGGCCGGGGCCGGG |

| | |
|---|---|
| | GCCGGGGCCGGGGCCGGGGCCGGGGCCGGGGCCGGGGCCGGGGCCGGGGCCGGGG<br>CCGGGGCCGGGGCCGGGGCCGGGGCCGGGGCCGGGGCCGGGGCCGGGGCCGGGGC<br>CGGGGCCGGGGCCGGGGCCGGGGCCGGGGCCGGGGCCGGGGCCUCUCGAGUCUAG<br>AGGGCCGCAAGCCGGGAUCAUUCAUGGCAA |
| 5×Corn | GGUCGGCAGCACAUAUACUAGUCGACGGCCAUCAGUCGCCGGUCCCAAGCCCGGAUA<br>AAAUGGGAGGGGGCGGAAACCGCCUAACCAUGCCGACUGAUGGCAGUACCGUAGGC<br>GAAUUCUUGCCAUGUGUAUGUGGGAGGCCCGGAUAGCUCAGUCGGUAGAGCAGCGGC<br>CGCGCGAUUCGAGGAAGGAGGUCUGAGGAGGUCACUGAAUCGCGCGGCCGCGGGUCC<br>AGGGUUCAAGUCCCUGUUCGGGCCUCCCACAUACUCUGAUGAUCCAAAACUGCCCGAG<br>GAUCAUUCAUGGCAAACUCGACGUUCGUCAGACUCCUCGCGAUGUCUAUGCGGGAGG<br>CCCGGAUAGCUCAGUCGGUAGAGCAGCGGCCGCGCGAUUCGAGGAAGGAGGUCUGAG<br>GAGGUCACUGAAUCGCGCGGCCGCGGGUCCAGGGUUCAAGUCCCUGUUCGGGCCUCC<br>CGCAUAGUCUGAUCAUCCAAAACUGCCCGAGGGGAUGAUUCAUCGCGAAUCGCGACGA<br>UCGUCACUACCGGUUCACUUUGUAGACCAGGCUGGUGUACAACUUGCGACUUUGCCA<br>UGUGUAUGUGGGAGGCCCGGAUAGCUCAGUCGGUAGAGCAGCGGCCGCGCGAUUCGA<br>GGAAGGAGGUCUGAGGAGGUCACUGAAUCGCGCGGCCGCGGGUCCAGGGUUCAAGUC<br>CCUGUUCGGGCCUCCCACAUACUCUGAUGAUCCACGCGUAACAUGAGGAUCACCCAUG<br>UCGAGGUACCAUCAUUCAUGGCAAAGUCGCAAGUAUCGGUCAACCUAGUGGGCCCAGU<br>GACGAUCGUCGCCCAUCGCCUUGUGUCUGUUGGAGGCCCGGAUAGCUCAGUCGGUAG<br>AGCAGCGGCCGCGCGAUUCGAGGAAGGAGGUCUGAGGAGGUCACUGAAUCGCGCGGC<br>CGCGGGUCCAGGGUUCAAGUCCCUGUUCGGGCCUCCAACAGACUCUGACGAUCCAAA<br>ACUGCCCGAGGGGAUCGUUCAAGGCGAGAGCUCGUCUGACGAACGUCGAACUCGCCG<br>UGUGUGUGUAGGAGGCCCGGAUAGCUCAGUCGGUAGAGCAGCGGCCGCGCGAUUCGA<br>GGAAGGAGGUCUGAGGAGGUCACUGAAUCGCGCGGCCGCGGGUCCAGGGUUCAAGUC<br>CCUGUUCGGGCCUCCUACACACUCUGACGAUCCAAAACUGCCCGAGGGGAUCGUUCAC<br>GGCGAACUCACGUCGGUGGAGCUAGCGCCUACGGUACUGCCAUCAGUCGGCGUGGAC<br>UGUAGAACACUGCCAAUGCCGGUCCCAAGCCCGGAUAAAAGUGGAGGGUACAGUCCAC<br>GCUCUAGAGCGGACUUCGGUCCGCUUUUUACUAGGACCUGCAGGCAUGCAAGCUUGA<br>CGUCGGUUACCGAUAUCCAUAUGGCGACCGC |
| 5×Beetroot (with fluorogenic Pepper inserted) | GGUCGGCAGCACAUAUACUAGUCGACGGCCAUCAGUCGCCGGUCCCAAGCCCGGAUA<br>AAAUGGGAGGGGGCGGAAACCGCCUAACCAUGCCGACUGAUGGCAGUACCGUAGGC<br>GAAUUCUUGCCAUGUGUAUGUGGGAGGCCCGGAUAGCUCAGUCGGUAGAGCAGCGGC<br>CGGUUAGGCAGAGGUGGGUGGUGUGGAGGAGUAUCUGUCCGGCCGCGGGUCCAGGG<br>UUCAAGUCCCUGUUCGGGCCUCCCACAUACUCUGAUGAUCCAAAACUGCCCGAGGAUC<br>AUUCAUGGCAAACUCGACGUUCGUCAGACUCCUCGCGAUGUCUAUGCGGGAGGCCCG<br>GAUAGCUCAGUCGGUAGAGCAGCGGCCGGUUAGGCAGAGGUGGGUGGUGUGGAGGA<br>GUAUCUGUCCGGCCGCGGGUCCAGGGUUCAAGUCCCUGUUCGGGCCUCCCGCAUAGU<br>CUGAUCAUCCAAAACUGCCCGAGGGGAUGAUUCAUCGCGAAUCGCGACGAUCGUCACU<br>ACCGGUGACUCGCUAUCGGUUAUGGCAUGUACAACUUGCGACUUUGCCAUGUGUAUG<br>UGGGAGGCCCGGAUAGCUCAGUCGGUAGAGCAGCGGCCGGUUAGGCAGAGGUGGGU<br>GGUGUGGAGGAGUAUCUGUCCGGCCGCGGGUCCAGGGUUCAAGUCCCUGUUCGGGC<br>CUCCCACAUACUCUGAUGAUCCACGCGUGUAGGAGGGAUCCCCAAUCGUGGCGUGUC<br>GGCCUGCUUCGGCAGGCACUGGCGCCGGGAUCCCUCCUACGGUACCAUCAUUCAUGG<br>CAAAGUCGCAAGUAUCGGUCAACCUAGUGGGCCCAGUGACGAUCGUCGCCCAUCGCC<br>UUGUGUCUGUUGGAGGCCCGGAUAGCUCAGUCGGUAGAGCAGCGGCCGGUUAGGCAG<br>AGGUGGGUGGUGUGGAGGAGUAUCUGUCCGGCCGCGGGUCCAGGGUUCAAGUCCCU<br>GUUCGGGCCUCCAACAGACUCUGACGAUCCAAAACUGCCCGAGGGGAUCGUUCAAGG<br>CGAGAGCUCGUCUGACGAACGUCGAACUCGCCGUGUGUGUGUAGGAGGCCCGGAUAG<br>CUCAGUCGGUAGAGCAGCGGCCGGUUAGGCAGAGGUGGGUGGUGUGGAGGAGUAUC |

| | |
|---|---|
| | UGUCCGGCCGCGGGUCCAGGGUUCAAGUCCCUGUUCGGGCCUCCUACACACUCUGAC<br>GAUCCAAAACUGCCCGAGGGGAUCGUUCACGGCGAACUCACGUCGGUGGAGCUAGCG<br>CCUACGGUACUGCCAUCAGUCGGCGUGGACUGUAGAACACUGCCAAUGCCGGUCCCA<br>AGCCCGGAUAAAAGUGGAGGGUACAGUCCACGCUCUAGAGCGGACUUCGGUCCGCUU<br>UUUACUAGGACCUGCAGGCAUGCAAGCUUGACGUCGGUUACCGAUAUCCAUAUGGCGA<br>CCGC |
| Broc-NS | GGCACAGUGCUAUGAGUGUCGCGACGGAGACGGUCGGGUCCAGAUAGGCCAGUCGAC<br>AAGGUCUAUCUGUCGAGUAGAGUGUGGGCUCCGUCGCGUGCACAUUAGAGUCGCUGU<br>AUGCCACAGUCGACAAGUGGCGUACAGCGGCUCUAGUGUGCUGCACAGUGUCUGUGC<br>GACUGCACCCAGUCGACAAGGGUGUAGUCGCAUAGACAUUGUGCUCACUCGUAGCAUU<br>GUGCCUGUCUCCAGUCGACAAGGAGAUAG |
| Pep-NS | GGCACAGUGCUAUGAGUGUGCACGGGAUCCCCAAUCGUGGCGUGUCGGCCUGCCGCA<br>UCGCGAAAGUGGCAGGCACUGGCGCCGGGAUCCUGUGCUGCACAUUAGAGUCGCUGU<br>AUGACCCAUCGCGAAAGGGUCGUACAGCGGCUCUAGUGUGCUCGCGUGCCUCAGAGG<br>ACCUGUCACCAUCGCGAAAGGUGAUAGGUCCUUUGAGGUACGCGUCACUCGUAGCAU<br>UGUGCCUGUCUCCAUCGCGAAAGGAGAUAG |
| c2dB-47×CAG | GGCCGCACUCGCCGGUCCCAAGCCCGGAUAAAAUGGGAGGGGGCGGGAAACCGCCUA<br>ACCAUGCCGAGUGCGGCCGCUACCGUAGGCUCGCCGUGUAAUACGACUCACUAUAGG<br>GCAGCACAUAUACUAGUCGACUUGCCAUGUGUAUGUGGGAGACGGUCGGGUCCAUCU<br>GAGACGGUCGGGUCCAGAUAUUCGUAUCUGUCGAGUAGAGUGUGGGCUCAGAUGUCG<br>AGUAGAGUGUGGGCUCCCACAUACUCUGAUGAUCCAGACGGUCGGGUCCAUCUGAGA<br>CGGUCGGGUCCAGAUAUUCGUAUCUGUCGAGUAGAGUGUGGGCUCAGAUGUCGAGUA<br>GAGUGUGGGCUGGAUCAUUCAUGGCAAUCUAGAGAAUUCGGUCUCACAGCAGCAGCA<br>GCAGCAGCAGCAGCAGCAGCAGCAGCAGCAGCAGCAGCAGCAGCAGCAGCAGCAG<br>CAGCAGCAGCAGCAGCAGCAGCAGCAGCAGCAGCAGCAGCAGCAGCAGCAGCAGCAGC<br>AGCAGCAGCAGCAGUUUUUUUCCGCGGCUAGCAUAACCCCUUGCACGGCGAGCCUAC<br>GGUAGUGGCCGCGGUCGGCGUGGACUGUAGAACACUGCCAAUGCCGGUCCCAAGCCC<br>GGAUAAAAGUGGAGGGUACAGUCCACGC |
| cBroc-47×CUG | GGCCGCACUCGCCGGUCCCAAGCCCGGAUAAAAUGGGAGGGGGCGGGAAACCGCCUA<br>ACCAUGCCGAGUGCGGCCGCUUGCCAUGUGUAUCGGGGAGACGGUCGGGUCCAGAUA<br>UUCGUAUCUGUCGAGUAGAGUGUGGGCUCCCCGAUACUCUGAUGAUCCUCUAGAGAA<br>UUCCUGCUGCUGCUGCUGCUGCUGCUGCUGCUGCUGCUGCUGCUGCUGCUGCUGCU<br>GCUGCUGCUGCUGCUGCUGCUGCUGCUGCUGCUGCUGCUGCUGCUGCUGCUGCUGC<br>UGCUGCUGCUGCUGCUGCUGCUGCUGCUGCUGGCUAGCAUAAGGAUCAUUCAUGGCA<br>AGUGGCCGCGGUCGGCGUGGACUGUAGAACACUGCCAAUGCCGGUCCCAAGCCCGGA<br>UAAAAGUGGAGGGUACAGUCCACGC |
| cDNB-17×$G_4C_2$ | GGCCGCACUCGCCGGUCCCAAGCCCGGAUAAAAUGGGAGGGGGCGGGAAACCGCCUA<br>ACCAUGCCGAGUGCGGCCGCUACCGUAGGCUUGCCAUGUGUAUCGGGUGCCUUAUUC<br>CGGACGCCGGGCCCGAAUGCUGCUACGGCAGUCGAAGACAACAUCGCGCCCUUCGGA<br>GGCACCCGAUACUCUGAUGAUCCUUGAAUUCGCUAGCGGGGCCGGGGCCGGGGCCGG<br>GGCCGGGGCCGGGGCCGGGGCCGGGGCCGGGGCCGGGGCCGGGGCCGGGGCCGGG<br>GCCGGGGCCGGGGCCGGGGCCGGGGCCUCUCGAGAUGCGUCGGGAUCAUUCAUGGC<br>AAGCCUACGGUAGUGGCCGCGGUCGGCGUGGACUGUAGAACACUGCCAAUGCCGGUC<br>CCAAGCCCGGAUAAAAGUGGAGGGUACAGUCCACGC |
| cNS3 | GGCCGCACUCGCCGGUCCCAAGCCCGGAUAAAAUGGGAGGGGGCGGGAAACCGCCUA<br>ACCAUGCCGAGUGCGGCCGCUUGCCAUGUGUAUCGCACAGUGCUAUGAGUGUCGCGA<br>CGGAGACGGUCGGGUCCAGAUAGGCCAGUCGACAAGGUCUAUCUGUCGAGUAGAGUG<br>UGGGCUCCGUCGCGUGCACAUUAGAGUCGCUGUAUGCCACAGUCGACAAGUGGCGUA<br>CAGCGGCUCUAGUGUGCUGCACAGUGUCUGUGCGACUGCACCCAGUCGACAAGGGUG |

| | |
|---|---|
| | UAGUCGCAUAGACAUUGUGCUCACUCGUAGCAUUGUGCGAUACUCUGAUGAUCCUUCG<br>GGAUCAUUCAUGGCAAGUGGCCGCGGUCGGCGUGGACUGUAGAACACUGCCAAUGCC<br>GGUCCCAAGCCCGGAUAAAAGUGGAGGGUACAGUCCACGC |
| cNS2 | GGCCGCACUCGCCGGUCCCAAGCCCGGAUAAAAUGGGAGGGGGCGGGAAACCGCCUA<br>ACCAUGCCGAGUGCGGCCGCUUGCCAUGUGUAUCGCACAGUGCUAUGAGUGUCGCGA<br>CGGAGACGGUCGGGUCCAGAUAGGCCAGUCGACAAGGUCUAUCUGUCGAGUAGAGUG<br>UGGGCUCCGUCGCGUGCACAUUAGAGUCGCUGUAUGCCACAGUCGACAAGUGGCGUA<br>CAGCGGCUCUAGUGUGCUGCACAGUGUCUGUGCGACUGCACCCAACGAACAAGGGUG<br>UAGUCGCAUAGACAUUGUGCUCACUCGUAGCAUUGUGCGAUACUCUGAUGAUCCUUCG<br>GGAUCAUUCAUGGCAAGUGGCCGCGGUCGGCGUGGACUGUAGAACACUGCCAAUGCC<br>GGUCCCAAGCCCGGAUAAAAGUGGAGGGUACAGUCCACGC |
| cNS0 | GGCCGCACUCGCCGGUCCCAAGCCCGGAUAAAAUGGGAGGGGGCGGGAAACCGCCUA<br>ACCAUGCCGAGUGCGGCCGCUUGCCAUGUGUAUCGCACAGUGCUAUGAGUGUCGCGA<br>CGGAGACGGUCGGGUCCAGAUAGGCCAUGGCGAAAGGUCUAUCUGUCGAGUAGAGUG<br>UGGGCUCCGUCGCGUGCACAUUAGAGUCGCUGUAUGCCACAUAGCACAAGUGGCGUA<br>CAGCGGCUCUAGUGUGCUGCACAGUGUCUGUGCGACUGCACCCAACGAACAAGGGUG<br>UAGUCGCAUAGACAUUGUGCUCACUCGUAGCAUUGUGCGAUACUCUGAUGAUCCUUCG<br>GGAUCAUUCAUGGCAAGUGGCCGCGGUCGGCGUGGACUGUAGAACACUGCCAAUGCC<br>GGUCCCAAGCCCGGAUAAAAGUGGAGGGUACAGUCCACGC |
| ctNS2 | GGCCGCACUCGCCGGUCCCAAGCCCGGAUAAAAUGGGAGGGGGCGGGAAACCGCCUA<br>ACCAUGCCGAGUGCGGCCGCUUGCCAUGUGUAUCGCACAGUGCUAUGAGUGUCGCGA<br>CGGAGACGGUCGGGUCCAGAUAGGCCAGUCGACAAGGUCUAUCUGUCGAGUAGAGUG<br>UGGGCUCCGUCGCGUGCACAUUAGAGUCGCUGUAUGCCACAGUCGACAAGUGGCGUA<br>CAGCGGCUCUAGUGUGCUCACUCGUAGCAUUGUGCGAUACUCUGAUGAUCCUUCGGG<br>AUCAUUCAUGGCAAGUGGCCGCGGUCGGCGUGGACUGUAGAACACUGCCAAUGCCGG<br>UCCCAAGCCCGGAUAAAAGUGGAGGGUACAGUCCACGC |
| csNS3 | GGCCGCACUCGCCGGUCCCAAGCCCGGAUAAAAUGGGAGGGGGCGGGAAACCGCCUA<br>ACCAUGCCGAGUGCGGCCGCUUGCCAUGUGUAUCUGAGUGUCGCGAGAGACGGUCGG<br>GUCCAGAUAGGCCAGUCGACAAGGUCUAUCUGUCGAGUAGAGUGUGGGCUCUCGCGU<br>GCACAUGCCACAGUCGACAAGUGGCGUGUGCUGCACAGCACCCAGUCGACAAGGGUG<br>UUGUGCUCACUCGGAUACUCUGAUGAUCCUUCGGGAUCAUUCAUGGCAAGUGGCCGC<br>GGUCGGCGUGGACUGUAGAACACUGCCAAUGCCGGUCCCAAGCCCGGAUAAAAGUGG<br>AGGGUACAGUCCACGC |
| csNS0 | GGCCGCACUCGCCGGUCCCAAGCCCGGAUAAAAUGGGAGGGGGCGGGAAACCGCCUA<br>ACCAUGCCGAGUGCGGCCGCUUGCCAUGUGUAUCUGAGUGUCGCGAGAGACGGUCGG<br>GUCCAGAUAGGCCAUGGCGAAAGGUCUAUCUGUCGAGUAGAGUGUGGGCUCUCGCGU<br>GCACAUGCCACAUAGCACAAGUGGCGUGUGCUGCACAGCACCCAACGAACAAGGGUGU<br>UGUGCUCACUCGGAUACUCUGAUGAUCCUUCGGGAUCAUUCAUGGCAAGUGGCCGCG<br>GUCGGCGUGGACUGUAGAACACUGCCAAUGCCGGUCCCAAGCCCGGAUAAAAGUGGA<br>GGGUACAGUCCACGC |
| cuNS3 | GGCCGCACUCGCCGGUCCCAAGCCCGGAUAAAAUGGGAGGGGGCGGGAAACCGCCUA<br>ACCAUGCCGAGUGCGGCCGCUUGCCAUGUGUAUCGCACAGUGCUAUAGAAAAAAAGAC<br>GGAGACGGUCGGGUCCAGAUAGGCCAGUCGACAAGGUCUAUCUGUCGAGUAGAGUGU<br>GGGCUCCGGAAGAAAAAAAUUAGAGUCGCUGUAUGCCACAGUCGACAAGUGGCGUACA<br>GCGGCUCUAGGAAAAAAGAAGGUGUCUGUGCGACUGCACCCAGUCGACAAGGGUGUA<br>GUCGCAUAGACAUAAAAAAAAAAAGUAGCAUUGUGCGAUACUCUGAUGAUCCUUCGGG<br>AUCAUUCAUGGCAAGUGGCCGCGGUCGGCGUGGACUGUAGAACACUGCCAAUGCCGG<br>UCCCAAGCCCGGAUAAAAGUGGAGGGUACAGUCCACGC |

| | |
|---|---|
| cuNS0 | GGCCGCACUCGCCGGUCCCAAGCCCGGAUAAAAUGGGAGGGGGCGGGAAACCGCCUA ACCAUGCCGAGUGCGGCCGCUUGCCAUGUGUAUCGCACAGUGCUAUAGAAAAAAAGAC GGAGACGGUCGGGUCCAGAUAGGCCAUGGCGAAAGGUCUAUCUGUCGAGUAGAGUGU GGGCUCCGGAAGAAAAAAAUUAGAGUCGCUGUAUGCCACAUAGCACAAGUGGCGUACA GCGGCUCUAGGAAAAAAGAAGGUGUCUGUGCGACUGCACCCAACGAACAAGGGUGUAG UCGCAUAGACAUAAAAAAAAAAAGUAGCAUUGUGCGAUACUCUGAUGAUCCUUCGGGA UCAUUCAUGGCAAGUGGCCGCGGUCGGCGUGGACUGUAGAACACUGCCAAUGCCGGU CCCAAGCCCGGAUAAAAGUGGAGGGUACAGUCCACGC |
| cY2 | GGCCGCACUCGCCGGUCCCAAGCCCGGAUAAAAUGGGAGGGGGCGGGAAACCGCCUA ACCAUGCCGAGUGCGGCCGCUUGCCAUGUGUAUCGGGAGACGGUCGGGUCCAGAUAG GCCAGUCGACAAGGUCUAUCUGUCGAGUAGAGUGUGGGCUCCCGAUACUCUGAUGAU CCUCUGUGCGACUGCACCCAGUCGACAAGGGUGUAGUCGCAUAGAGGAUCAUUCAUG GCAAGUGGCCGCGGUCGGCGUGGACUGUAGAACACUGCCAAUGCCGGUCCCAAGCCC GGAUAAAAGUGGAGGGUACAGUCCACGC |
| cY1 | GGCCGCACUCGCCGGUCCCAAGCCCGGAUAAAAUGGGAGGGGGCGGGAAACCGCCUA ACCAUGCCGAGUGCGGCCGCUUGCCAUGUGUAUCGGGAGACGGUCGGGUCCAGAUAG GCCAGUCGACAAGGUCUAUCUGUCGAGUAGAGUGUGGGCUCCCGAUACUCUGAUGAU CCUCUGUGCGACUGCACCCAACGAACAAGGGUGUAGUCGCAUAGAGGAUCAUUCAUGG CAAGUGGCCGCGGUCGGCGUGGACUGUAGAACACUGCCAAUGCCGGUCCCAAGCCCG GAUAAAAGUGGAGGGUACAGUCCACGC |
| c0 | GGCCGCACUCGCCGGUCCCAAGCCCGGAUAAAAUGGGAGGGGGCGGGAAACCGCCUA ACCAUGCCGAGUGCGGCCGCUUGCCAUGUGUAUCGGGGAGACGGUCGGGUCCAGAUA UUCGUAUCUGUCGAGUAGAGUGUGGGCUCCCCGAUACUCUGAUGAUCCUUCGGGAUC AUUCAUGGCAAGUGGCCGCGGUCGGCGUGGACUGUAGAACACUGCCAAUGCCGGUCC CAAGCCCGGAUAAAAGUGGAGGGUACAGUCCACGC |
| P-cNS3 | GGCCGCACUCGCCGGUCCCAAGCCCGGAUAAAAUGGGAGGGGGCGGGAAACCGCCUA ACCAUGCCGAGUGCGGCCGCUUGCCAUGUGUAUCGCACAGUGCUAUGAGUGUGCACG GGAUCCCCAAUCGUGGCGUGUCGGCCUGCCGCAUCGCGAAAGUGGCAGGCACUGGCG CCGGGAUCCUGUGCUGCACAUUAGAGUCGCUGUAUGACCCAUCGCGAAAGGGUCGUA CAGCGGCUCUAGUGUGCUCGCGUGCCUCAGAGGACCUGUCACCAUCGCGAAAGGUGA UAGGUCCUUUGAGGUACGCGUCACUCGUAGCAUUGUGCGAUACUCUGAUGAUCCUUC GGGAUCAUUCAUGGCAAGUGGCCGCGGUCGGCGUGGACUGUAGAACACUGCCAAUGC CGGUCCCAAGCCCGGAUAAAAGUGGAGGGUACAGUCCACGC |
| BP-cNS3 | GGCCGCACUCGCCGGUCCCAAGCCCGGAUAAAAUGGGAGGGGGCGGGAAACCGCCUA ACCAUGCCGAGUGCGGCCGCUUGCCAUGUGUAUCGCACAGUGCUAUGAGUGUCGCGA CGGAGACGGUCGGGUCCAGAUAGGCCAGGUACCAAGGUCUAUCUGUCGAGUAGAGUG UGGGCUCCGUCGCGUGCACAUUAGAGUCGCUGUAUGCCACAGGUACCAAGUGGCGUA CAGCGGCUCUAGUGUGCUGCACGGAUCCCCAAUCGUGGCGUGUCGGCCUGCCGCAGG UACCAAGUGGCAGGCACUGGCGCCGGGAUCCUGUGCUCACUCGUAGCAUUGUGCGAU ACUCUGAUGAUCCUUCGGGAUCAUUCAUGGCAAGUGGCCGCGGUCGGCGUGGACUGU AGAACACUGCCAAUGCCGGUCCCAAGCCCGGAUAAAAGUGGAGGGUACAGUCCACGC |
| rP-cNS3 | GGCCGCACUCGCCGGUCCCAAGCCCGGAUAAAAUGGGAGGGGGCGGGAAACCGCCUA ACCAUGCCGAGUGCGGCCGCUUGCCAUGUGUAUCGGUGUUACGAUGCUCACUGCGCU GGGAUCCCCAAUCGUGGCGUGUCGGCCUGCCUGCACAGCUGAAGCGGGCAGGCACUG GCGCCGGGAUCCCAGCGCUCGUGUGAUCUCGGCGACAUGCGGUCACAGCUGAAGACC GUAUGUCGCUGAGAUUACACGUCGUGUUACAGAUACGCUGAUGUGGCACAGCUGAAG CCACGUCAGCGUGUCUGUGACACGUGUGAGUAUCGUGACACCGAUACUCUGAUGAUC CUUCGGGAUCAUUCAUGGCAAGUGGCCGCGGUCGGCGUGGACUGUAGAACACUGCCA AUGCCGGUCCCAAGCCCGGAUAAAAGUGGAGGGUACAGUCCACGC |

| | |
|---|---|
| P-cNS3-MS2 | GGCCGCACUCGCCGGUCCCAAGCCCGGAUAAAAUGGGAGGGGGCGGGAAACCGCCUA<br>ACCAUGCCGAGUGCGGCCGCUUGCCAUGUGUAUCGCACAGUGCUAUGAGUGUGCACG<br>GGAUCCCCAAUCGUGGCGUGUCGGCCUGCCGCAUCGCGAAAGUGGCAGGCACUGGCG<br>CCGGGAUCCUGUGCUGCACAUUAGAGUCGCUGUAUGACCCAUCGCGAAAGGGUCGUA<br>CAGCGGCUCUAGUGUGCUCGCGUGCCUCAGAGGACCUGUCACCAUCGCGAAAGGUGA<br>UAGGUCCUUUGAGGUACGCGUCACUCGUAGCAUUGUGCGAUACUCUGAUGAUCCACG<br>CGUAACAUGAGGAUCACCCAUGUCGAGGUACCAUGGAUCAUUCAUGGCAAGUGGCCGC<br>GGUCGGCGUGGACUGUAGAACACUGCCAAUGCCGGUCCCAAGCCCGGAUAAAAGUGG<br>AGGGUACAGUCCACGC |
| cNS3-Pepper (TAR v2) | GGCCGCACUCGCCGGUCCCAAGCCCGGAUAAAAUGGGAGGGGGCGGGAAACCGCCUA<br>ACCAUGCCGAGUGCGGCCGCUUGCCAUGUGUAUCGCACAGUGCUAUGAGUGUCGCGA<br>CGGAGACGGUCGGGUCCAGAUAGGCCAGUCGACAAGGUCUAUCUGUCGAGUAGAGUG<br>UGGGCUCCGUCGCGUGCACAUUAGAGUCGCUGUAUGCCACAGUCGACAAGUGGCGUA<br>CAGCGGCUCUAGUGUGCUGCACAGUGUCUGUGCGACUGCACCCAGUCGACAAGGGUG<br>UAGUCGCAUAGACAUUGUGCUCACUCGUAGCAUUGUGCGAUACUCUGAUGAUCCGCUA<br>GCAAAGGCUCGUUGAGCUCAUUAGCUCCGAGCCCGAGGUACCGGAUCAUUCAUGGCA<br>AGUGGCCGCGGUCGGCGUGGACUGUAGAACACUGCCAAUGCCGGUCCCAAGCCCGGA<br>UAAAAGUGGAGGGUACAGUCCACGC |
| cPepper (TAR v2) | GGCCGCACUCGCCGGUCCCAAGCCCGGAUAAAAUGGGAGGGGGCGGGAAACCGCCUA<br>ACCAUGCCGAGUGCGGCCGCUUGCCAUGUGUAUGUGGGACGCGUUGCCACGUUUCCC<br>ACAUACUCUGAUGAUCCGCUAGCAAAGGCUCGUUGAGCUCAUUAGCUCCGAGCCCGAG<br>GUACCGGAUCAUUCAUGGCAAGCGGCCGCGGUCGGCGUGGACUGUAGAACACUGCCA<br>AUGCCGGUCCCAAGCCCGGAUAAAAGUGGAGGGUACAGUCCACGC |
| P-cNS3-coB | GGCCGCACUCGCCGGUCCCAAGCCCGGAUAAAAUGGGAGGGGGCGGGAAACCGCCUA<br>ACCAUGCCGAGUGCGGCCGCUUGCCAUGUGUAUCGCACAGUGCUAUGAGUGUGCACG<br>GGAUCCCCAAUCGUGGCGUGUCGGCCUGCCGCAUCGCGAAAGUGGCAGGCACUGGCG<br>CCGGGAUCCUGUGCUGCACAUUAGAGUCGCUGUAUGACCCAUCGCGAAAGGGUCGUA<br>CAGCGGCUCUAGUGUGCUCGCGUGCCUCAGAGGACCUGUCACCAUCGCGAAAGGUGA<br>UAGGUCCUUUGAGGUACGCGUCACUCGUAGCAUUGUGCGAUACUCUGAUGAUCCUAG<br>UCGUAUUAAUUUCGAUAAGCCAGUAAGCAGUGGGAUGGAUCAUUCAUGGCAAGUGGCC<br>GCGGUCGGCGUGGACUGUAGAACACUGCCAAUGCCGGUCCCAAGCCCGGAUAAAAGU<br>GGAGGGUACAGUCCACGC |
| cNS3-Sw | GGCCGCACUCGCCGGUCCCAAGCCCGGAUAAAAUGGGAGGGGGCGGGAAACCGCCUA<br>ACCAUGCCGAGUGCGGCCGCUUGCCAUGUGUAUCGCACAGUGCUAUGAGUGUCGCGA<br>CGGAGACGGUCGGGUCCAGAUAGGCCAGUCGACAAGGUCUAUCUGUCGAGUAGAGUG<br>UGGGCUCCGUCGCGUGCACAUUAGAGUCGCUGUAUGCCACAGUCGACAAGUGGCGUA<br>CAGCGGCUCUAGUGUGCUGCACAGUGUCUGUGCCAUGUGUUCUAGUCGACAAAGAAC<br>UCUGAUGAAUAACCGAAGUAGUUUAUGGGCUACCGAAAUCAUUCAUGGCAUAGACAUU<br>GUGCUCACUCGUAGCAUUGUGCGAUACUCUGAUGAUCCUUCGGGAUCAUUCAUGGCA<br>AGUGGCCGCGGUCGGCGUGGACUGUAGAACACUGCCAAUGCCGGUCCCAAGCCCGGA<br>UAAAAGUGGAGGGUACAGUCCACGC |
| cY2-Sw | GGCCGCACUCGCCGGUCCCAAGCCCGGAUAAAAUGGGAGGGGGCGGGAAACCGCCUA<br>ACCAUGCCGAGUGCGGCCGCUUGCCAUGUGUAUCGGGAGACGGUCGGGUCCAGAUAG<br>GCCAGUCGACAAGGUCUAUCUGUCGAGUAGAGUGUGGGCUCCCGAUACUCUGAUGAU<br>CCUCUGUGCCAUGUGUUCUAGUCGACAAAGAACUCUGAUGAAUAACCGAAGUAGUUUA<br>UGGGCUACCGAAAUCAUUCAUGGCAUAGAGGAUCAUUCAUGGCAAGUGGCCGCGGUC<br>GGCGUGGACUGUAGAACACUGCCAAUGCCGGUCCCAAGCCCGGAUAAAAGUGGAGGG<br>UACAGUCCACGC |

### SUPPLEMENTARY FIGURES

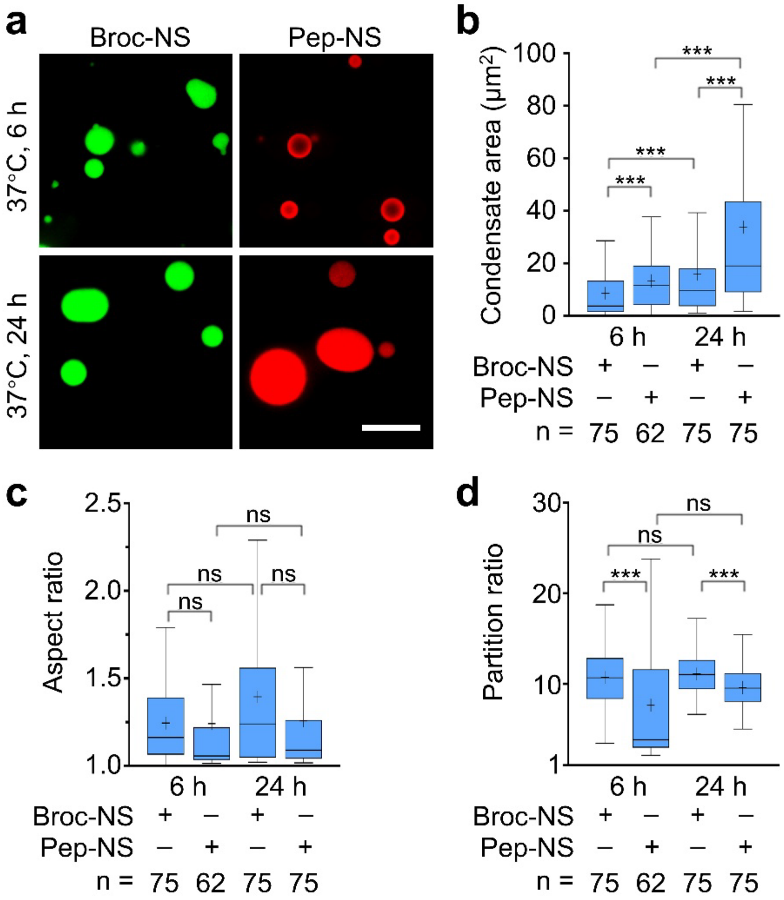


**Fig. S1 | *In vitro* condensate formation of RNA nanostar scaffolds using a "melt-and-hold" protocol.**

**a,** Confocal fluorescence images show RNA nanostar condensates formed after the "melt-and-hold" procedure (70 °C heat, rapid cooling to 37 °C, and incubated at 37 °C for 6 or 24 h). Two nanostar variants were tested: the "A-type" and "B-type", in which the Pepper aptamer replaces the malachite green aptamer in A-type, while Broccoli remains embedded in B-type, yielding Pep-NS and Broc-NS, respectively. Imaging was performed at room temperature in solutions containing 4 μM RNA, 20 mM $MgCl_2$, and either 80 μM DFHBI-1T or 2uM HBC620. Scale bar: 10 μm. The vesicle-like morphology observed in Pep-NS at 6 h may reflect percolated structures or limited dye accessibility that result in preferential periphery fluorescence.

**b–d,** Quantification of condensate area, aspect ratio, and partition ratio—defined as the mean fluorescence intensity inside condensate relative to background—for each scaffold type from **a** and displayed as Tukey box-and-whisker plots. Boxes represent the 25th and 75th percentiles; central lines indicate the medians; whiskers extend to 1.5× the interquartile range; means are indicated by "+". Data represent n condensates from ≥3 independent replicates. Statistical significance was assessed by two-way ANOVA with Tukey's post-hoc test (****p* ≤ 0.001; ns, *p* > 0.05).

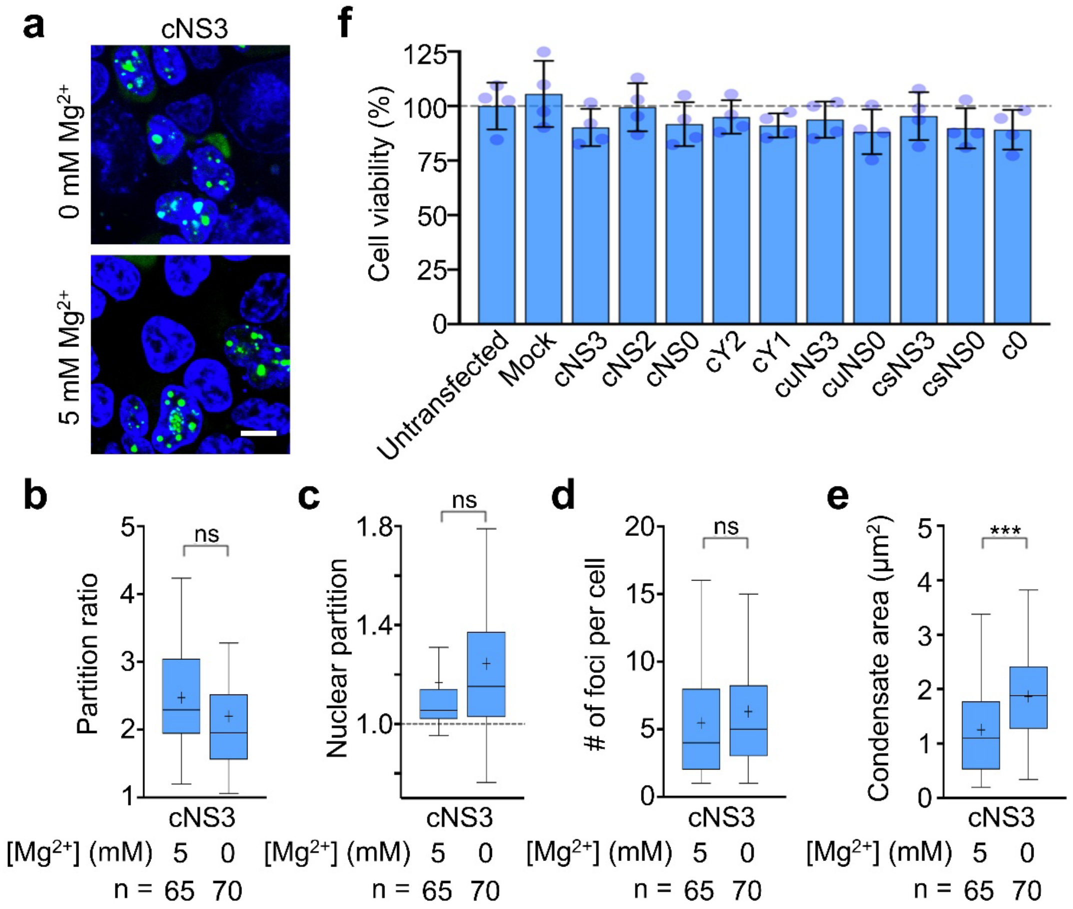


**Fig. S2 | Cytotoxicity and effect of magnesium ions on cellular nanostar condensates.**

**a,** Confocal fluorescence images of HEK293T cells expressing the circularized RNA nanostar scaffold cNS3 via the pAVU6+27-Tornado vector. Live-cell imaging was performed after a 30-min incubation with 40 μM DFHBI-1T and either 0 or 5 mM $Mg^{2+}$. Hoechst 33342 was used for nuclear staining. Scale bar: 10 μm.

**b–e,** Quantification of partition ratio, condensate area, condensate number per cell, and nuclear partition for each condition in **a**. Data are represented as Tukey box-and-whisker plots, where boxes represent the 25th and 75th percentiles, central lines indicate the medians, whiskers extend to 1.5× the interquartile range, and means are indicated by "+". A total of n cells per scaffold type were analyzed from ≥3 independent replicates. Statistical significance was determined using two-tailed Student's *t*-tests relative to cNS3 (***$p \leq 0.001$; ns, $p > 0.05$).

**f,** Cell proliferation determined by XTT assay after a 24-hour incubation at 37°C following transfection with plasmids encoding various RNA nanostar scaffolds. Untransfected and Fugene mock-transfected cells served as controls. Cell viability was normalized to untransfected cells. Shown are the mean ± standard deviation values from four independent replicates. Compared to untransfected cells, no significant differences ($p > 0.05$) were observed through two-tailed student's *t*-tests.

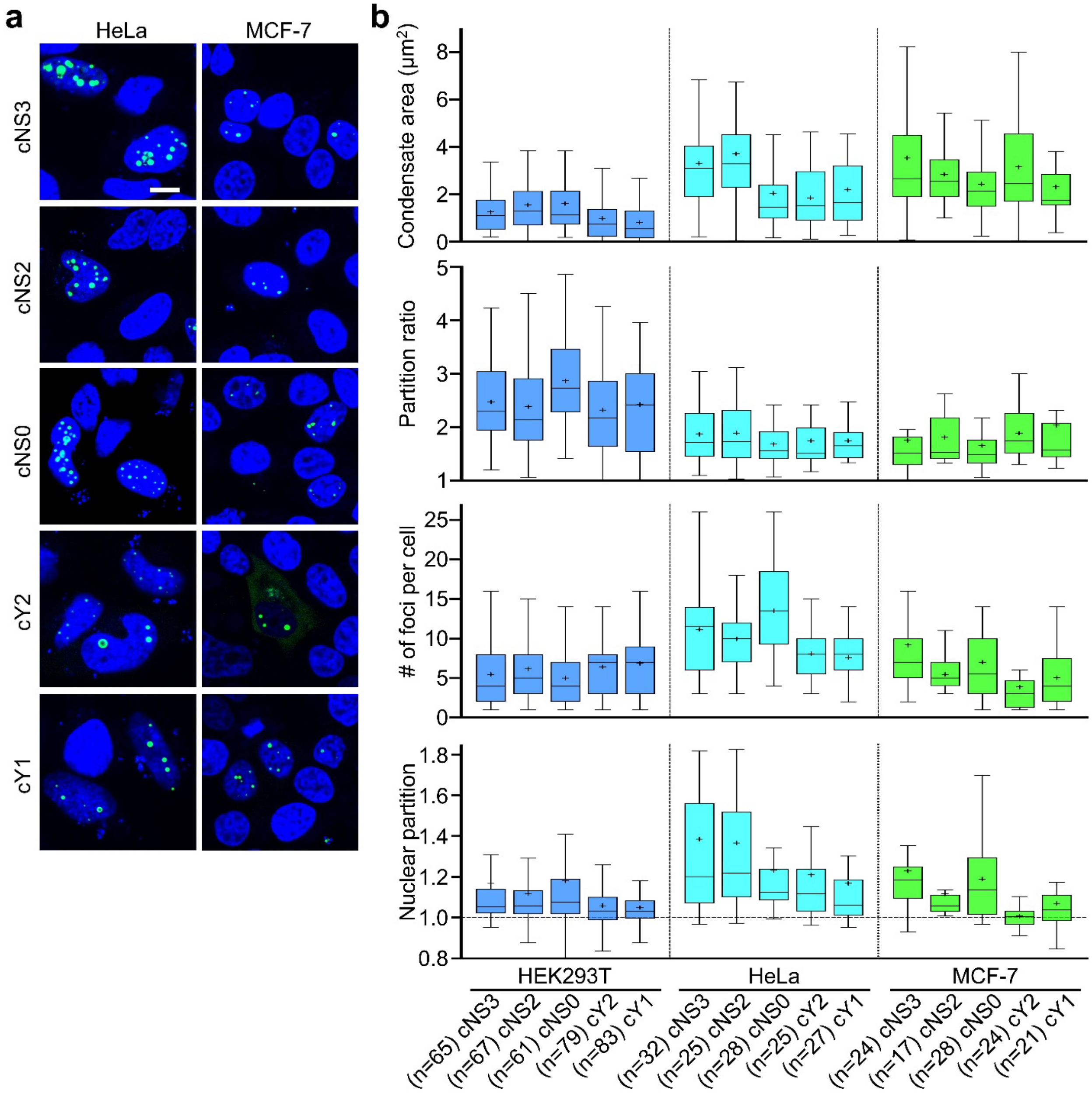


**Fig. S3 | RNA nanostar scaffolds induce condensate formation across diverse cell lines.**

**a,** Confocal fluorescence images of HeLa and MCF-7 cells expressing circularized RNA nanostar scaffolds (cNS3, cNS2, cNS0, cY2 and cY1) via the pAVU6+27-Tornado vector. Live-cell imaging was performed after a 30-min incubation with 40 μM DFHBI-1T and 5 mM $Mg^{2+}$. Hoechst 33342 was used for nuclear staining. Scale bar: 10 μm.

**b,** Quantification of partition ratio, condensate area, condensate number per cell, and nuclear partition for each scaffold in HEK293T, HeLa, and MCF-7 cells. Data are represented as Tukey box-and-whisker plots, where boxes represent the 25th and 75th percentiles, central lines indicate the medians, whiskers extend to 1.5× the interquartile range, and means are indicated by “+”. A total of n cells per scaffold type were analyzed from at least two independent replicates.

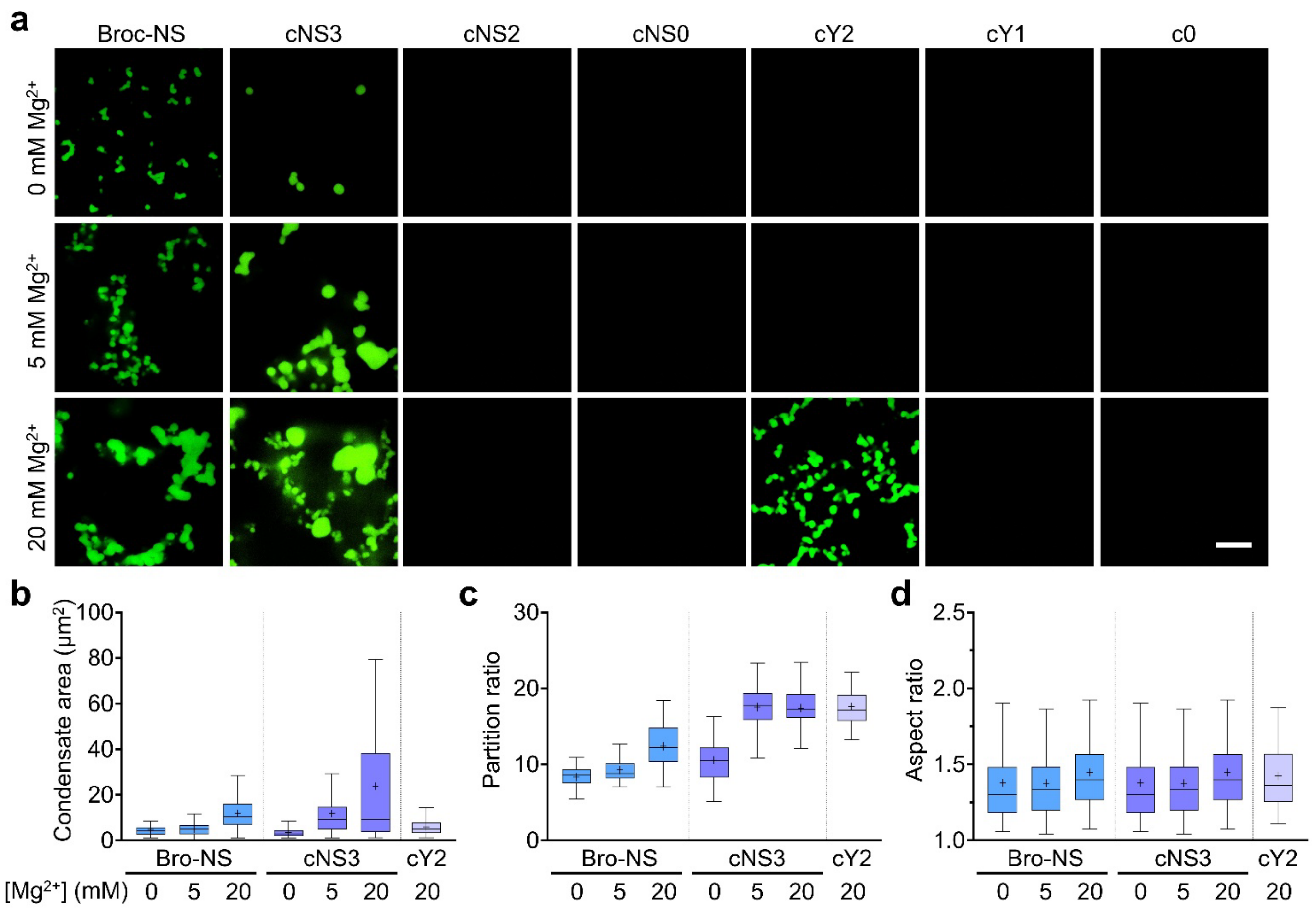


**Fig. S4 | Effect of magnesium ions and kissing loops on RNA nanostar condensate formation *in vitro.***

**a,** Confocal fluorescence images of RNA nanostar condensates formed using the "melt-and-hold" protocol (70 °C heating, rapid cooling to 37 °C, and a 12-h incubation at 37 °C). The "B-type" nanostar (Broc-NS) contains an embedded Broccoli aptamer and four active kissing loops. Linearized variants of additional scaffolds (cNS3, cNS2, cNS0, cY2, cY1 and c0), each differing in kissing-loop and stem number, were also examined. Imaging was performed at room temperature in a buffer containing 4 μM RNA, 80 μM DFHBI-1T, 40 mM HEPES, 100 mM KCl, 500 mM NaCl, and varying $MgCl_2$ concentrations. Scale bar: 10 μm.

**b,** Quantification of condensate area, aspect ratio, and partition ratio for each condensate-forming condition from **a**. Data are represented as Tukey box-and-whisker plots, where boxes represent the 25th and 75th percentiles, central lines indicate the medians, whiskers extend to 1.5× the interquartile range, and means are indicated by "+". 50 condensates per condition were analyzed from at least two independent replicates.

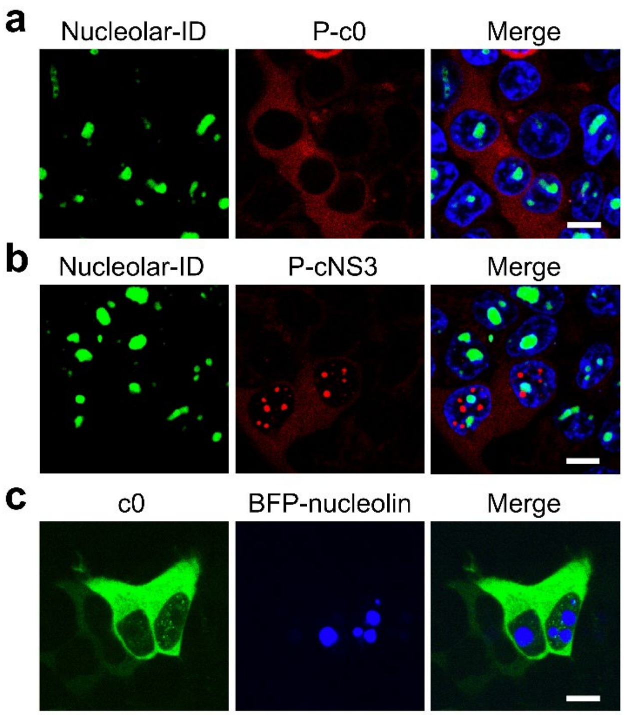


**Fig. S5 | Cellular nanostar condensates do not colocalize with nucleoli.**

**a**, **b,** Confocal fluorescence images of HEK293T cells stained with Nucleolar-ID, which labels nucleoli (green). Cells express either the Pepper-fused circularized F30 (P-c0) or the Pepper-embedded nanostar (P-cNS3). P-c0 lacks a nanostar structure and kissing loops, while P-cNS3 contains three active kissing loops. Live-cell imaging was performed after a 30-min incubation with 5 μM HBC620 and 5 mM $Mg^{2+}$. Hoechst 33342 was used for nuclear staining. Scale bar: 10 μm.

**c,** Confocal fluorescence images of HEK293T cells co-expressing BFP-nucleolin and either Broccoli-tagged circularized F30 (c0) or Broccoli-embedded nanostar (cNS3). Nucleolin is the major nucleolar component. Live-cell imaging was performed after a 30-min incubation with 40 μM DFHBI-1T and 5 mM $Mg^{2+}$. Scale bar: 10 μm.

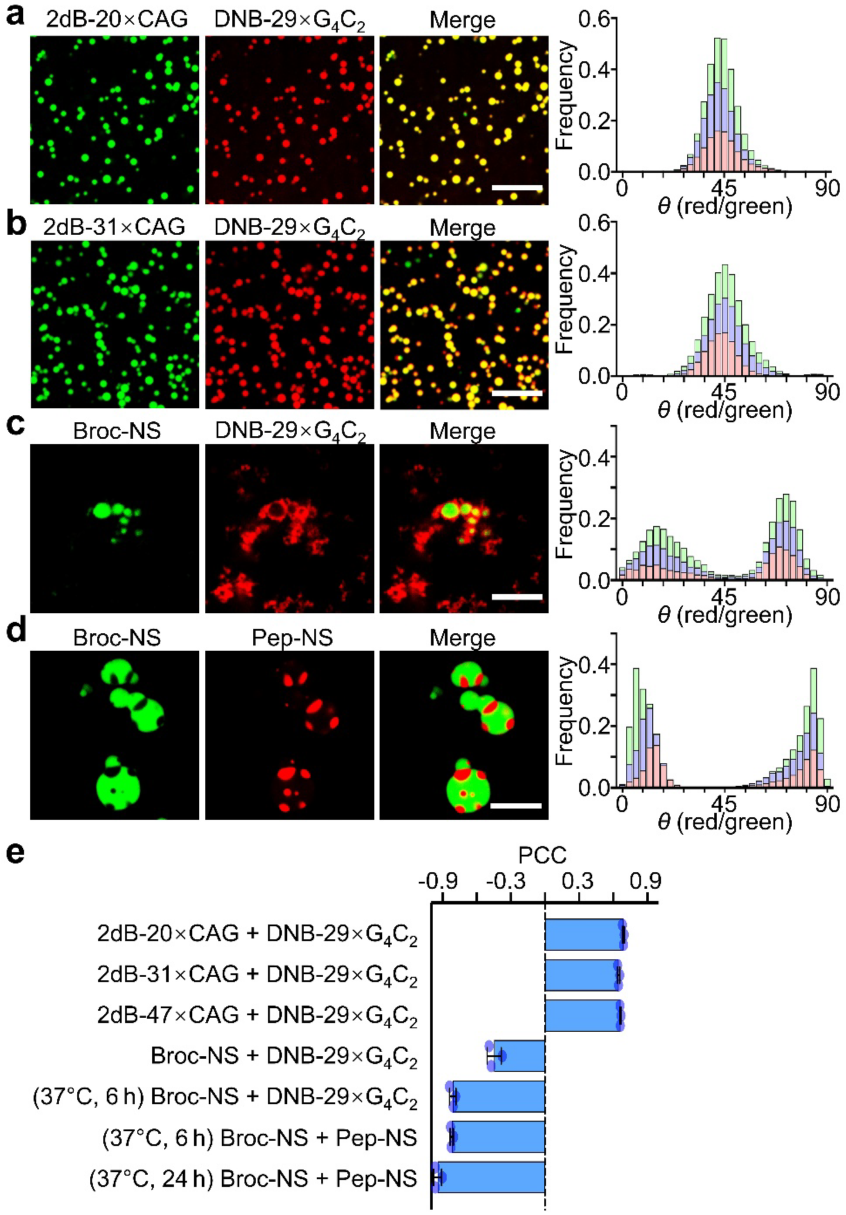


**Fig. S6 | Orthogonality of RNA condensates *in vitro* among different RNA scaffolds.**

**a**, **b,** Confocal fluorescence images showing condensate formation and miscibility between pairs of RNA scaffolds. Fluorogenic 2d×Broccoli (2dB) fused to 20× or 31×CAG repeats and DNB fused to 29×$G_4C_2$ (2 μM each) were mixed and imaged after annealing (95 °C for 2 min, slow cooling –0.5 °C/min to room temperature). Imaging buffers contained 20 mM $MgCl_2$, 80 μM DFHBI-1T, and 0.5 μM TMR-DN. Orthogonality was assessed by coordinate-angle mapping of pixel-wise intensity ratios between the DNB and Broccoli channels; a single peak near 45° indicates high overlap. Histograms represent three independent replicates.

**c**, **d,** Confocal fluorescence images of Broccoli-embedded nanostar (Broc-NS) paired with either DNB-29×$G_4C_2$ or Pepper-embedded nanostar (Pep-NS) (2 μM each) following "melt-and-hold" incubation (70 °C for 10 min, rapid cooling to 37 °C, and holding for 6 h for Broc-NS/DNB-29×$G_4C_2$ or 24 h for Broc-NS/Pep-NS). Imaging buffers contained 20 mM $MgCl_2$, 80 μM DFHBI-1T, and either 0.5 μM TMR-DN or 2 μM HBC620. Orthogonality was assessed by coordinate angle mapping of pixel-wise intensity ratios between the red (DNB or Pepper) and green (Broccoli) channels; a single peak near 45° indicates high overlap, whereas distinct peaks near 0° and 90° indicate strong orthogonality. Histograms represent three independent replicates.

**e,** Pearson correlation coefficients (PCCs) calculated from three independent replicates for each scaffold pair. Positive PCC values (>0.6) indicate strong channel correlation and low orthogonality, values near zero indicate minimal correlation, and negative PCC values (<–0.6) indicates anticorrelated signals consistent with high orthogonality. Data are shown as Fisher Z-transformed mean ± SD.

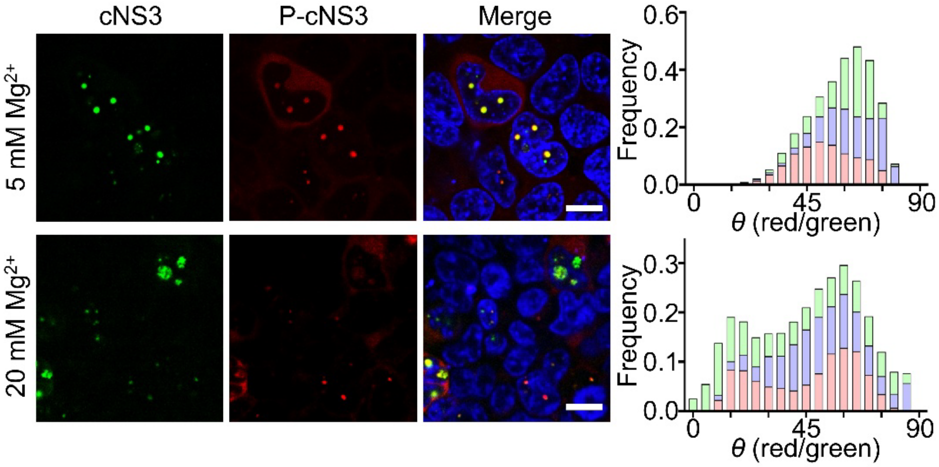


**Fig. S7 | Cellular RNA nanostar condensates show limited and $Mg^{2+}$-promoted orthogonality.**

Confocal fluorescence images of HEK293T cells co-transfected with the circular Broccoli-embedded nanostar (cNS3) and the circular Pepper-embedded nanostar (P-cNS3). Live-cell imaging was performed after a 30-min incubation with 40 μM DFHBI-1T, 5 μM HBC620 and either 5 mM or 20 mM $Mg^{2+}$. Hoechst 33342 was used for nuclear staining. Scale bar: 10 μm. Right panel: coordinate-angle mapping was used to quantify condensate orthogonality. Pixel-wise intensity ratios between the Pepper and Broccoli channels were converted to arctangent angles (*θ*). A single peak near 45° indicates substantial channel overlap and poor orthogonality, whereas distinct peaks near 0° and 90° indicate minimal overlap and strong orthogonality. Stacking histograms represent three independent replicates.

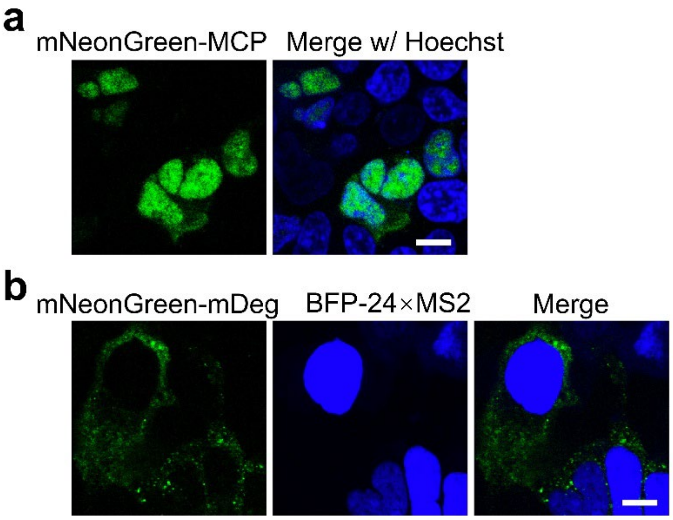


**Fig. S8 | Condensate-free cellular distributions of mNeonGreen-MCP and mNeonGreen-mDeg.**

**a,** Confocal fluorescence images of HEK293T cells expressing mNeonGreen-MCP. mNeonGreen-MCP contains a nuclear localization signal (NLS), resulting in homogenous nuclear fluorescence. Live-cell imaging was performed after a 30-min incubation with 40 μM DFHBI-1T and 5 mM $Mg^{2+}$. Hoechst 33342 was used for nuclear staining. Scale bar: 10 μm.

**b,** Confocal fluorescence images of HEK293T cells expressing BFP-24×MS2 and mNeonGreen-mDeg. The BFP-24×MS2 mRNA contains a 5'-UTR binding domain and 24 MS2 hairpins for visualization, producing a nucleus-localized BFP signal. mNeonGreen-mDeg is a degron-fused mNeonGreen-MCP variant that becomes stabilized only upon binding MS2-tagged mRNA, forming cytoplasmic foci. Live-cell imaging was performed after a 30-min incubation with 40 μM DFHBI-1T and 5 mM $Mg^{2+}$. Scale bar: 10 μm.

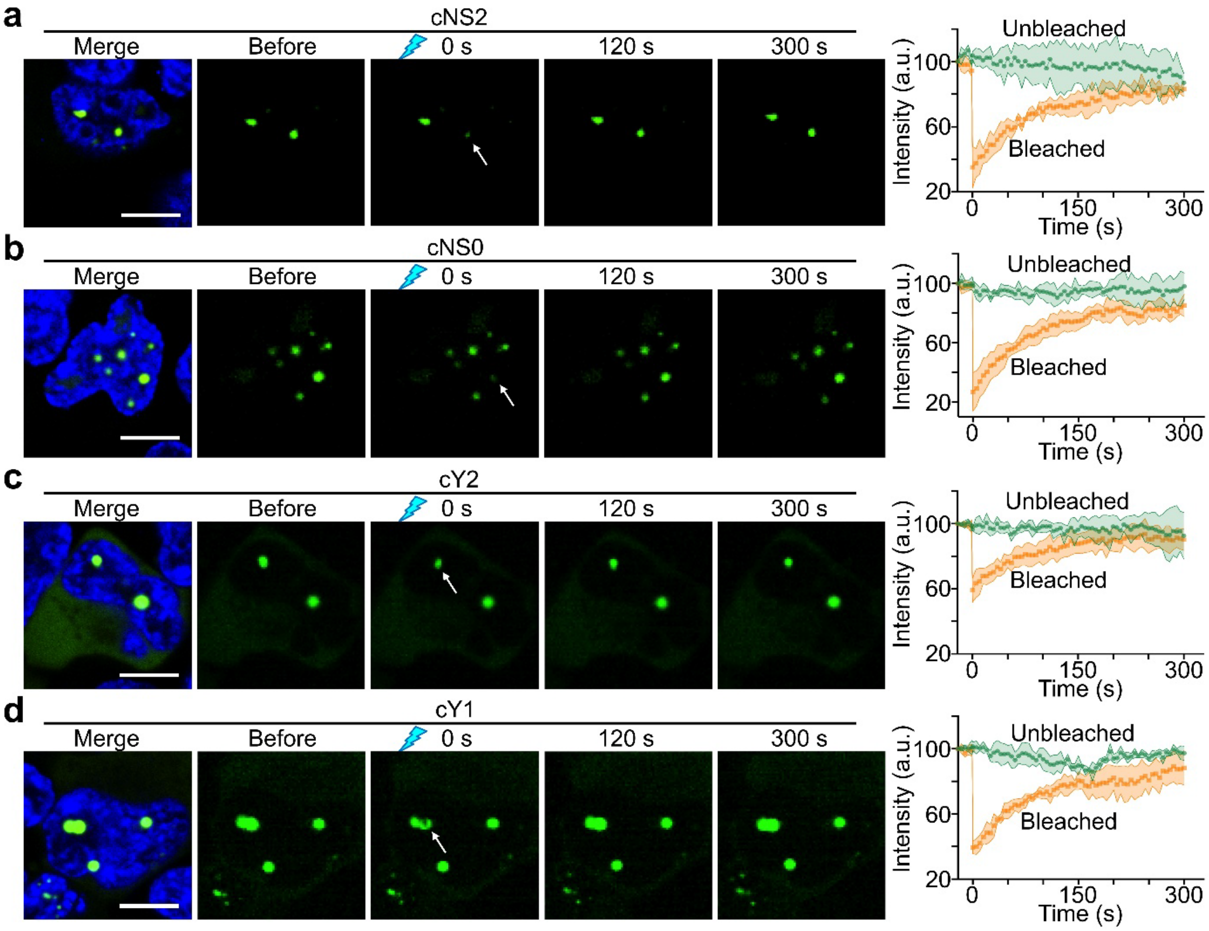


**Fig. S9 | Fluorescence recovery after photobleaching (FRAP) of RNA nanostar condensates in cells.**

**a–d,** Cellular FRAP analysis of HEK293T cells expressing circular cNS2, cNS0, cY2 or cY1 condensates. The Broccoli dye DFHBI-1T was used to monitor fluorescence recovery following photobleaching. Live-cell imaging was performed after a 30-min incubation with 40 μM DFHBI-1T and 5 mM $Mg^{2+}$. Hoechst 33342 was used for nuclear staining. The white arrow marks the photobleaching region. Right panels show FRAP trajectories of photobleached condensates (orange) and non-photobleached condensates (green). All condensate types exhibited rapid recovery with ~80% of initial fluorescence within ~200 s, indicating fast mobility and exchange of small-molecule DFHBI-1T within the condensates. Data represent mean ± SD from three independent replicates. Scale bar: 10 μm.

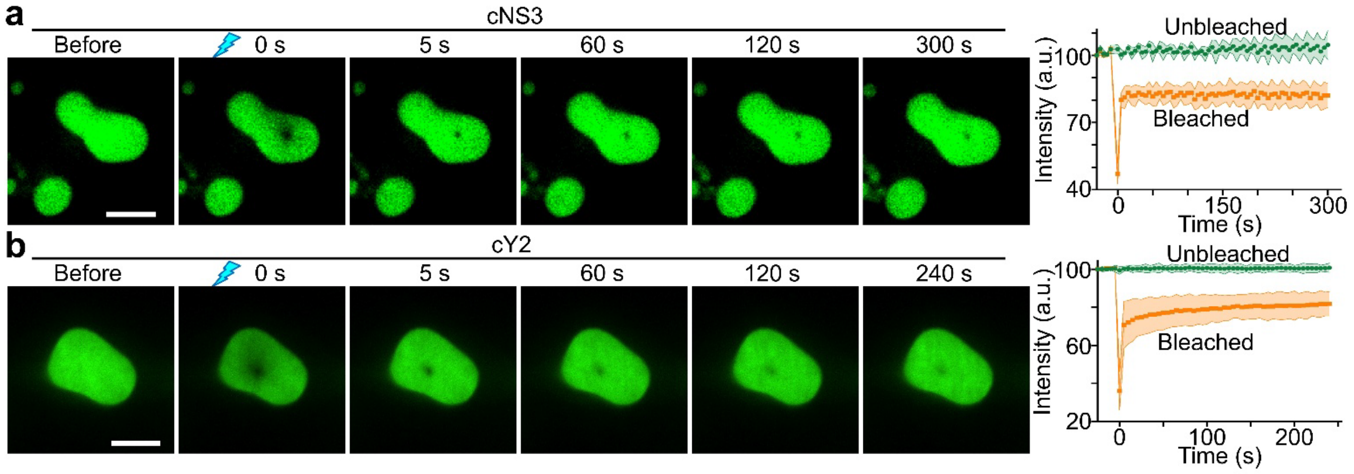


**Fig. S10 | FRAP of RNA nanostar condensates *in vitro*.**

**a**, **b,** FRAP analysis of *in vitro* condensates formed by linearized cNS3 and cY2 scaffolds labeled with th Broccoli dye DFHBI-1T. Condensates were generated in solutions containing 4 μM RNA, 80 μM DFHBI-1T, 40 mM HEPES, 100 mM KCl, 500 mM NaCl, and 20 mM $MgCl_2$ using the "melt-and-hold" protocol (70 °C for 10 min, rapid cooling to 37 °C, incubation at 37 °C for 12 h, then cooling to room temperature). Right panels show FRAP trajectories of photobleached condensates (orange) and non-photobleached condensates (green). Both scaffold types exhibited rapid recovery—70–80% of the initial signals within ~5 s—leaving an immobile fraction of ~20%. These results indicate fast dye diffusion within the condensates, with the possibility of a compact core that limits dye penetration. Data represent mean ± SD from three independent replicates. Scale bar: 5 μm.

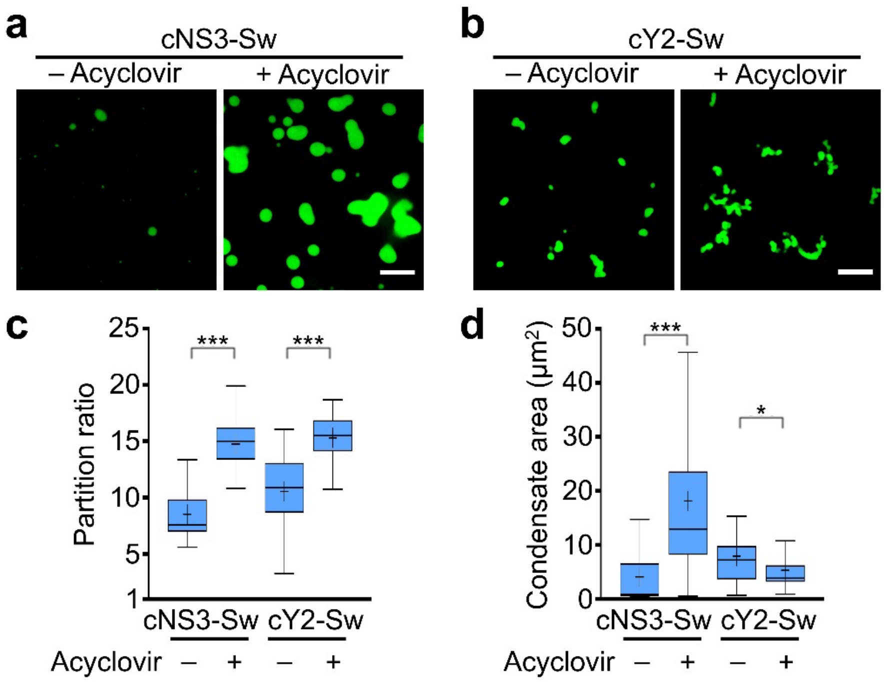


**Fig. S11 | Acyclovir-induced switchable RNA condensate formation *in vitro*.**

**a**, **b,** Acyclovir-induced condensate formation of switchable RNA scaffolds *in vitro.* The cNS3-Sw and cY2-Sw constructs incorporate an acyclovir-responsive RNA switch that disrupts one double-stranded RNA stem and kissing loop, therefore preventing condensate formation in the absence of ligand. Acyclovir binding restores the proper folding of RNA scaffolds and re-activates condensate formation. Linear cNS3-Sw and cY2-Sw were synthesized and subjected to the "melt-and-hold" protocol (70 °C heating, rapid cooling to 37 °C, and incubation at 37 °C for 24 h) in solutions containing 4 μM RNA and 20 mM $MgCl_2$. Vehicle (0.1% v/v DMSO) or 100 μM acyclovir was added prior to incubation. DFHBI-1T (80 μM) was added 30 min before imaging. Scale bar: 10 μm.

**c**, **d,** Quantification of partition ratio and condensate area. Data are represented as Tukey box-and-whisker plots, where boxes represent the 25th and 75th percentiles, central lines indicate the medians, whiskers extend to 1.5× the interquartile range, and means are indicated by "+". 50 condensates per condition were analyzed from at least two independent replicates. Statistical significance was assessed using two-tailed Student's *t*-tests (***$p \leq 0.001$; *$p \leq 0.05$).

### SUPPLEMENTARY VIDEOS

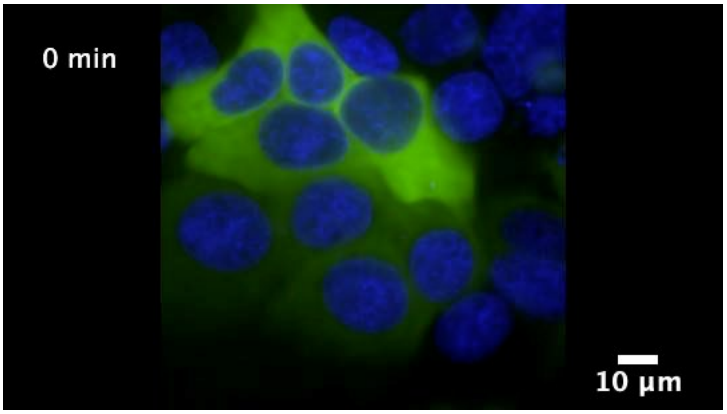


**Video S1a |** Fluorescence timelapse imaging of HEK293T cells expressing cY2-Sw immediately after addition of 100 µM acyclovir.

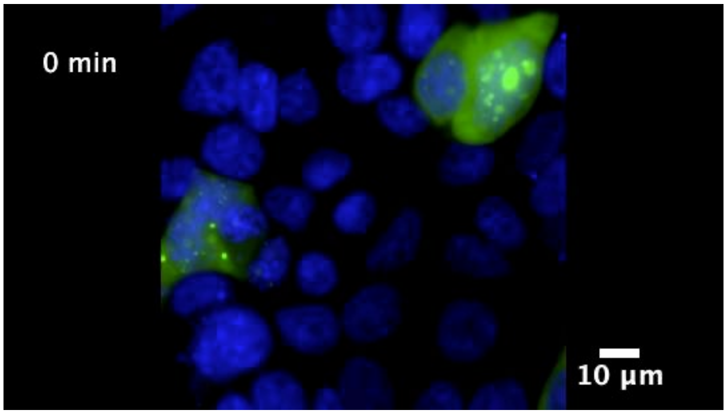


**Video S1b |** Fluorescence timelapse imaging of HEK293T cells expressing cY2-Sw after 4 h of incubation with 100 µM acyclovir followed by medium replacement.

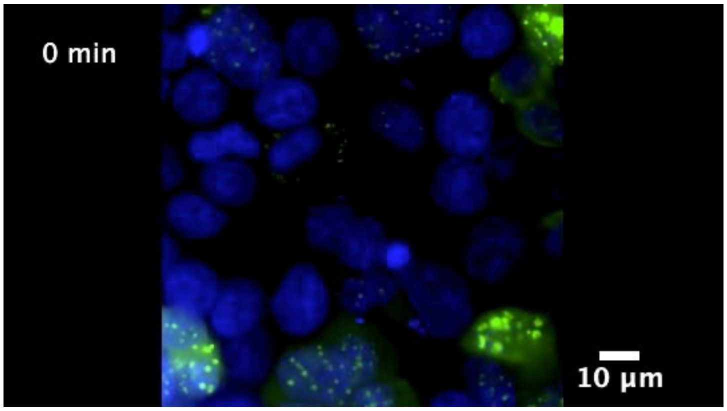


**Video S2a |** Fluorescence timelapse imaging of HEK293T cells expressing cNS3-Sw immediately after addition of 100 µM acyclovir.

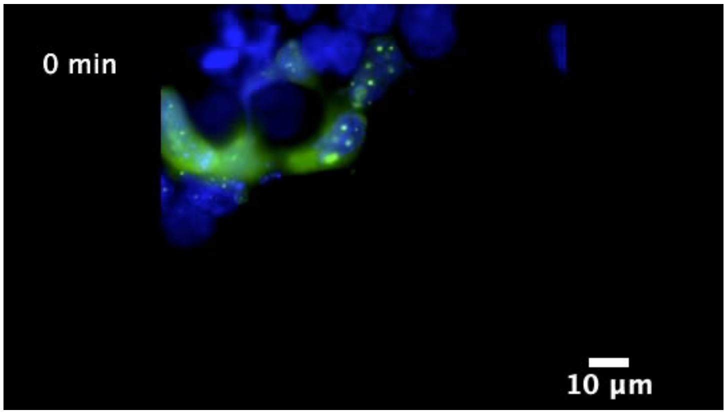


**Video S2b |** Fluorescence timelapse imaging of HEK293T cells expressing cNS3-Sw after 4 h of incubation with 100 µM acyclovir followed by medium replacement.
